# Prediction (or not) during language processing. A commentary on Nieuwland et al. (2017) and DeLong et al. (2005)

**DOI:** 10.1101/143750

**Authors:** Shaorong Yan, Gina R. Kuperberg, T. Florian Jaeger

**Affiliations:** Department of Brain and Cognitive Sciences, University of Rochester; Department of Psychology, Tufts University; Department of Psychiatry and the Athinoula A. Martinos Center for Biomedical Imaging, Massachusetts General Hospital, Harvard Medical School; Department of Brain and Cognitive Sciences, Department of Computer Science, Department of Linguistics, University of Rochester

**Keywords:** prediction, surprisal, Bayesian surprise, event-related potentials, hierarchical predictive processes, N400, N250

## Abstract

The extent to which language processing involves prediction of upcoming inputs remains a question of ongoing debate. One important data point comes from DeLong et al. (2005) who reported that an N400-like event-related potential correlated with a probabilistic index of upcoming input. This result is often cited as evidence for gradient probabilistic prediction of form and/or semantics, prior to the bottom-up input becoming available. However, a recent multi-lab study reports a failure to find these effects (Nieuwland et al., 2017). We review the evidence from both studies, including differences in the design and analysis approach between them. Building on over a decade of research on prediction since DeLong et al. (2005)’s original study, we also begin to spell out the computational nature of predictive processes that one might expect to correlate with ERPs that are evoked by a functional element whose form is dependent on an upcoming predicted word. For paradigms with this type of design, we propose an index of anticipatory processing, Bayesian surprise, and apply it to the updating of semantic predictions. We motivate this index both theoretically and empirically. We show that, for studies of the type discussed here, Bayesian surprise can be closely approximated by another, more easily estimated information theoretic index, the surprisal (or Shannon information) of the input. We re-analyze the data from Nieuwland and colleagues using surprisal rather than raw probabilities as an index of prediction. We find that surprisal is gradiently correlated with the amplitude of the N400, even in the data shared by Nieuwland and colleagues. Taken together, our review suggests that the evidence from both studies is compatible with anticipatory semantic processing. We do, however, emphasize the need for future studies to further clarify the nature and degree of form prediction, as well as its neural signatures, during language comprehension.

---

The role of prediction in language processing remains a topic of central interest to the field (Federmeier, 2007; Huettig & Mani, 2015; Kuperberg & Jaeger, 2016; Marta Kutas, DeLong, & Smith, 2011). Here we comment on a recent debate between Nieuwland and colleagues (Ito, Martin, & Nieuwland, 2016, 2017; Nieuwland et al., 2017) and DeLong and colleagues (DeLong, Urbach, & Kutas, 2005, 2017a,b).

An important study published in 2005 by DeLong et al. (2005) has been taken to provide an answer to three important questions about prediction. First, the study has been cited as evidence that predictions are generated prior to bottom-up evidence becoming available. Second, that predictions can be generated at both the level of semantic features and phonological form. And third, that these predictions are *probabilistic* in nature. There have been studies published prior to and after DeLong et al. reporting both behavioral (Allopenna, Magnuson, & Tanenhaus, 1998; Altmann & Kamide, 2007; Dahan & Tanenhaus, 2004) and neural signatures of predictive processing (Dikker & Pylkkänen, 2013; Otten, Nieuwland, & Van Berkum, 2007; Piai et al., 2016; Van Berkum, Brown, Zwitserlood, Kooijman, & Hagoort, 2005; Wicha, Moreno, & Kutas, 2004). However, DeLong et al. (2005) was one of the first studies that spoke to all three of these questions. This makes the results of a 9-lab study by Nieuwland et al. (2017), who recently report that they failed to replicate DeLong et al.’s (2005) findings, highly relevant to discussions of prediction in language processing. We use this debate as a welcome opportunity to review the two studies, and to speak to more general questions about the role of prediction in language comprehension.

We first describe the original DeLong et al. (2005) study. Then we describe the recent replication attempt by Nieuwland et al. (2017). We discuss important differences in the design and analysis of the two studies, and why they affect the conclusions that can be drawn from the failure to replicate. Following this summary, we discuss an alternative index of predictability to that used by both DeLong et al. (2005) and Nieuwland et al. (2017), motivated by recent research in psycho- and neurolinguistics. We find that this alternative index, *surprisal*, does, in fact, yield a significant effect for the data shared by Nieuwland et al. (2017).

We then consider the design of DeLong et al. (2005) in relation to three questions that we consider essential to any discussion of predictive processing (see Kuperberg & Jaeger, 2016). The first concerns the *timing* of prediction in relation to the appearance of the bottom-up input. Specifically, is activity at a given level of representation pre-activated ahead of new bottom-up information arriving or being decoded at that level of representation? The second concerns the level(s) of representation at which predictions are generated and updated (e.g. event structure, semantic features, form features etc.). The third concerns the candidate set of predicted entities. Specifically, is prediction *gradient*, that is, *probabilistically* conditioned on contextual expectations? Or is prediction an *all-or-nothing* phenomenon, entailing the prediction of just one or at most a very small set of candidates? Guided by these questions, we spell out different hypotheses about the predictive chain that could lead to effects such as those reported in DeLong et al. (2005) and similar studies (Van Berkum et al., 2005; Wicha et al., 2004). We formalize a probabilistic index of the hypothesized predictive processes in terms of Bayesian surprise, and compare it—both theoretically and empirically—to the index of predictability used in most ERP studies addressing this question thus far (cloze probabilities). Taken together, all these considerations lead us to conclude that is too early to dismiss the evidence for prediction observed in DeLong et al.’s original study.

## Background and summary of studies

### DeLong et al. (2005)

DeLong et al. (2005) measured event-related potentials (ERPs) in sentences like (1), in which the sentence context creates a range of constraints for a specific article+noun combination (‘a kite’). The words of interest then either confirm this expectation with the expected article+noun combination (‘a kite’), or disconfirm this expectation with a plausible yet unexpected continuation (e.g. ‘an airplane’). The authors focused on modulation of the N400—a negative going ERP component that peaks at around 400ms after stimulus onset, and that reflects the ease of semantically processing incoming information (for recent reviews, see Kuperberg, 2016; Kutas & Federmeier, 2011).

(1) The day was breezy so the boy went outside to fly a kite/an airplane ….

DeLong and colleagues carried out two main analyses in which they analyzed the relationship between the amplitude of the N400 and an estimate of the contextual predictability of both the article and the noun. In both analyses, contextual predictability was estimated using the cloze procedure (Taylor, 1953; more on the pros and cons of this approach below), and the N400 amplitude was operationalized as average activity between 200-500ms after the onset of words of interest. DeLong and colleagues sorted ERP trials by their cloze probability into 10 equally-sized bins (from 0-10% to 90-100% cloze). They then calculated the average ERP amplitude between 200-500ms within each bin in each participant at all 26 electrode sites, before averaging across participants.

They first analyzed the relationship between ERP amplitude evoked by the noun (e.g. “kite” in (1)) and the cloze probability of the noun. They found that, at centro-parietal electrode sites, the average amplitude of the waveform evoked by the noun in each bin inversely correlated with the average cloze probability of the noun in each bin. The centro-parietal scalp distribution of this effect was consistent with that of the N400. In other words, consistent with prior work (Kutas & Hillyard, 1984), the lower the cloze probability of the noun, the larger (more negative) the amplitude of the N400. This was interpreted as evidence for gradient probabilistic semantic processing at the point of the noun.

In addition, DeLong et al. examined the relationship between the amplitude of ERPs evoked by the preceding article, again averaged between 200-500ms, and the cloze probability of the article. As described below, their design allowed them to test whether comprehenders had pre-activated both the semantic and phonological features of the noun at the point the article was encountered. These were reasonable choices at the time of the paper’s original publication. However, as we discuss later, it is an open question whether the cloze probability of the article is the most appropriate measure for addressing these questions, or whether the full 200-500ms time window is the most sensitive time window to test hypotheses about the specific level of representation at which prediction takes place.

In DeLong et al.’s experiment, the expected target noun (e.g., ‘kite’ in (1)) was preceded by an indefinite article. Indeed, the target nouns in the experimental stimuli were *always* preceded by an indefinite article—either ‘a’ or ‘an’, depending on the phonology of the target noun (e.g., ‘a’ for ‘kite’ in (1)). Crucially, the unexpected noun always required the *opposite* indefinite article (e.g., ‘an’ for ‘airplane’ in (1)). How often the highly expected target noun required ‘a’ versus ‘an’ was counterbalanced across items and across participants. This design made it possible to address the question of whether comprehenders had generated not only semantic, but also form (phonological) predictions about the noun at the point at which the article was encountered.^1^ Specifically, the assumption was that any ERP modulation at an article that mismatches the phonological form of a predicted noun (e.g. ERP modulation on ‘an’ following a context like (1), which predicts the semantic features of <kite>), should only be seen if participants had actually predicted the form of ‘kite’ at this point. DeLong and colleagues carried out a similar analysis to the one that they carried out at the noun. Once again, they found evidence for a correlation, again at centro-parietal electrode sites, although they note that the topographic distribution of the effect was more right lateralized than at the noun (see Figure 1c, DeLong et al., 2005). The correlation observed on the article was numerically weaker than on the noun (though apparently statistically similar “at some electrode sites”, DeLong et al., 2005, p. 1119). DeLong et al. refer to this modulation as an N400 effect. They argued that these findings provide evidence that “anticipatory processing can happen not only for conceptual or semantic features but also for specific phonological word forms”, that “the system makes graded predictions” and that “articles, too, are predicted and integrated with context”.

**Figure 1.**
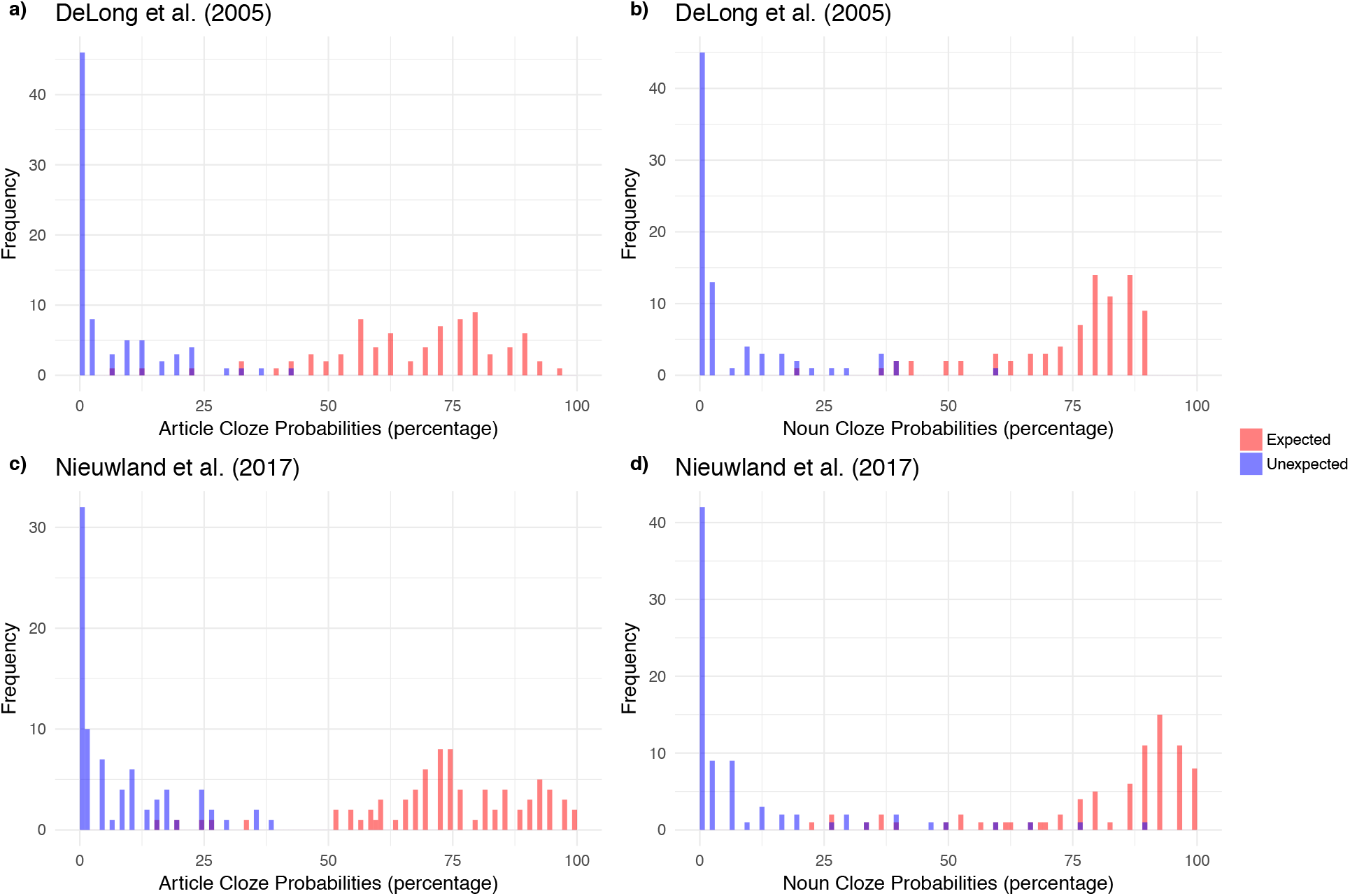
Histogram of cloze probabilities for articles (left panels) and nouns (right panels) for both expected (red) and unexpected (blue) items collected by both DeLong et al. (2005) (upper panels) and Nieuwland et al. (2017) (lower panels). We note that the leftmost bars contain not only items with 0 cloze probability, but also items with very small, but positive cloze probability.

### Nieuwland et al. (2017)

**Nieuwland et al. (2017)** present a large-scale replication attempt of DeLong et al. (2005), including 9 separate replications in 9 different labs across the United Kingdom. Their results on the article, they argue, call into question DeLong et al.’s conclusions that comprehenders probabilistically predict the form of the noun ahead of the article’s semantic features becoming available.

Whereas an earlier replication attempt by some of the same authors (Ito et al., 2016) followed DeLong et al.’s design relatively loosely, Nieuwland et al. (2017) aim for a close replication of both procedure and stimuli. We note, however, that a variety of different ERP recording systems were used, and that the nine replication experiments differed in other aspects both from each other and the original study, ranging from the numbers of participants in each individual study to methods of preprocessing and artifact rejection (see DeLong et al., 2017a for discussion of some of these issues). The stimuli are the same as in DeLong et al. (2005), with two exceptions. The first deviation is intended: Nieuwland and colleagues made minor modifications to the sentences to accommodate the differences in spelling, vocabulary, and cultural background between American English and British English. The second deviation from DeLong et al. is likely unintended and perhaps critical: while DeLong et al. (2005) included 116 fillers interspersed between 80 experimental stimuli (see DeLong et al., 2016), the 9-lab replication attempts by Nieuwland and colleagues only presented the 80 experimental stimuli. We return to this issue and its potential relevance below.

Nieuwland and colleagues reported two sets of analyses. The cloze probabilities (at both the noun and the article) that were used in all analyses were obtained from a new norming experiment with British participants, so as to accommodate potential differences in cloze probabilities between the (intended) populations. In addition, Nieuwland and colleagues employed exactly the same binning of cloze probabilities as DeLong and colleagues, and they also used the same time-windows over which the ERP signal was averaged (200-500ms).

In the first set of analyses—intended as a close replication of the correlation analyses performed by DeLong et al. (2005)—Nieuwland et al. (2017) examined the data separately for each of the nine labs. In each of these analyses, they used the same filters (.2 - 15Hz) for the EEG signal as used by DeLong et al., but they did not employ baseline correction. No such correction was mentioned in DeLong et al., 2005, 2017b, although, in fact, DeLong and colleagues used the same −500~0ms pre-article baseline for both the article and noun analyses (K. DeLong, p.c.).

In six of the labs, Nieuwland et al. (2017) report a significant correlation between the noun’s cloze probabilities and the N400 amplitude on the noun, with a similar classic N400 centro-parietal topographic distribution as that reported by DeLong et al. (2005); in two of the labs, the correlation trended, and in one lab it was insignificant (for further discussion of this, see DeLong et al., 2017a).

In contrast to the results on the noun, the correlation between the cloze probabilities on the article and N400 amplitude on the article failed to reach significance in any of the labs. In fact, most labs exhibited a numerical trend towards the opposite direction, i.e. a larger negativity in association with higher article cloze probabilities, which was significant in two of the labs. One lab did reveal a correlation between the negativity evoked between 200-500ms and cloze probability in the expected direction, but this was significant at left frontal electrodes rather than at central parietal sites as found by DeLong et al. (2005) (see Figure 1, Nieuwland et al., 2017).

In the second set of analyses, Nieuwland et al. (2017) go beyond the original analyses presented by DeLong et al., (2005) by analyzing the trial-level data. These analyses were conducted with linear mixed-effect regression, with cloze probability as a fixed-effect predictor and random by-participant and by-item intercepts as well as slopes for cloze probability. They did not include a random effect for lab because they argued that “there were only 9 laboratories, and laboratory was not a predictor of theoretical interest.” Instead, they treated lab as a fixed-effect predictor and also included the interaction between lab and cloze probabilities. They found that including lab as a predictor did not significant increase model fit, and so they only reported the model results without lab as a predictor. Additionally, for this single trial analysis, a few changes were made. Unlike for the individual lab analyses, Nieuwland and colleagues used unfiltered data and a baseline correction of −100~0ms. In addition, instead of running separate models for each electrode, they averaged the N400 amplitude across six central-parietal electrodes (Cz, C3, C4, Pz, P3, P4) and used this average as the dependent variable.

At the noun, this approach once again replicates the significant effect found by DeLong et al. (2005). At the article, however, Nieuwland et al. once again find that the cloze probability of the article does not statistically predict ERP modulation at the article, though we note that the effect is numerically in the predicted direction (*p* < .13).^2^ Nieuwland and colleagues argue that the numerical trend “at least partly reflected the effects of slow signal drift that existed before the articles were presented” and that it did not reflect ERP modulation induced by the articles themselves. They also present additional Bayesian analyses, arguing that these further support the absence of an N400 effect on the article. We describe these analyses in more detail below as part of our critique below.

Finally, to rule out the possibility that participants were just insensitive to the article manipulation, after each session, Nieuwland et al. (2017) ran a control experiment on the same participants (in each of the nine labs), in which the same critical nouns were preceded by either a correct or incorrect article (e.g. *an/*a apple*). Incorrect noun-article combination did elicit a larger P600, indicating that participants knew and were sensitive to the phonological constraint on the pre-nominal indefinite article. It is worth noting, however, that the nouns were embedded in sentences that were different from those used in the main experiment.

Based on these findings, Nieuwland et al. (2017) question the findings by DeLong and colleagues, and conclude that the claim “that prediction is probabilistic, rather than all-or-none, is now questionable”. They consider their findings a “challenge to the theory that comprehenders predict upcoming words, including their initial phonemes, through implicit production”.

## What can be inferred from this replication attempt?

Nieuwland and all the labs who took part in their study are to be applauded for dedicating the significant time and resources necessary for nine(!) replication attempts of an effect that is much cited and of high relevance to theory building in neurolinguistics. They further generously shared their preprocessed data, averaged across the 200-500ms time window, publically prior to publication, facilitating follow-up analyses by other researchers. Before we proceed, we also would also like to acknowledge the prompt and open responses to all of our follow-up questions by both Nieuwland and colleagues and DeLong and colleagues.

We begin by reviewing differences between the replication attempt and the original study. Second, we review the additional Bayesian analyses provided by Nieuwland et al. (2017) and conclude that they are less informative than might initially appear. Third, we turn our critique to three critical issues that are not unique to Nieuwland et al. (2017), but apply equally to other studies on prediction, including the original study by DeLong et al. (2005). All three of these final issues relate to the operationalization of predictability in terms of cloze probabilities. We also present one post-hoc analysis of the 9-lab-data shared by Nieuwland et al. (2017)—the only one we performed—that suggests that an alternative and theoretically-motivated index of probabilistic prediction, *surprisal*, actually reveals a significant effect in Nieuwland et al.’s datasets (even after correcting for multiple comparisons). After presenting our critique, we revisit the conclusions offered by Nieuwland and colleagues in the light of the three points outlined above that we consider to be critical in studying and interpreting probabilistic predictive effects in language comprehension.

## Differences from the original study

Due to omission of important methodological details in DeLong et al. (2005; as acknowledged by DeLong et al., 2017b), the 9-lab replication attempt by Nieuwland and colleagues differs from the original study in ways that are arguably important.

### No baseline correction used in the correlation analyses of the individual labs’ data

As noted above, in the correlation analyses carried out in the individual labs (on both the noun and the article), Nieuwland et al. (2017) did not use a baseline correction. This is in contrast to the analyses carried out by DeLong et al. (2005), who did use a baseline correction of −500-0ms (DeLong, p.c.), although, unfortunately, DeLong and colleagues did not report the use of this baseline in their original paper. We note this here because there is some evidence that this difference in analysis matters. As noted above, in the trial-level analysis for which Nieuwland et al. *did* use a baseline correction (−100-0ms on unfiltered data), there was a numerical trend of the data in the predicted direction—the same direction in which DeLong et al. (2005) report a significant effect.

To explore this further, we conducted an additional mixed-effects regression on the same trial-level data (−100-0ms baseline, unfiltered data). Besides cloze probability, we also included lab, as well as its interaction with cloze probability, as fixed-effect predictors (using the code that Nieuwland and colleagues shared online). We found that seven out of nine labs exhibit a trend in the *same* direction as in DeLong et al. (2005). In this re-analysis of Nieuwland and colleagues’ data, the main effect of cloze probability on the article was marginally significant (*p* < .13), in the same direction for which DeLong et al. (2005) found a significant effect (*p* < .05).

### Lack of fillers

All nine experiments presented in Nieuwland et al. (2017) lacked filler trials: participants saw 80 sentences with constraining contexts that were predictive of a specific noun (to various degrees, see Figure 1 below), and in half of the trials, this expectation was not met. This differs from the original experiment carried out by DeLong et al. (2005), which included 116 filler items.

It is possible that this difference between the two studies influenced modulation of the N400 on both the noun and the article. This is because repeated prediction mismatches might lead comprehenders to adapt what they predict, or even whether they predict at all. A number of behavioral studies have found that readers can adapt their expectations when they are repeatedly mismatched (e.g., Fine & Jaeger, 2016; Fine, Jaeger, Farmer, & Qian, 2013; Fraundorf & Jaeger, 2016; Kaschak, 2007; Kaschak & Glenberg, 2004)—adaptation that is captured by a Bayesian model of expectation adaptation (Fine et al., 2010; Kleinschmidt, Fine, & Jaeger, 2012). Some studies have found that even strong expectations can be changed drastically—based on the recent statistics of the input—within as few as 20 sentences (Farmer, Fine, Yan, Cheimariou, & Jaeger, 2014; Fine et al., 2013). Similarly fast expectation adaptation has also been observed at the lexical level (Brown-Schmidt, 2009; Creel, Aslin, & Tanenhaus, 2008; Yan & Farmer, 2015; see also Yildirim, Degen, Tanenhaus, & Jaeger, 2016).

Most relevant to the present study, there is evidence that the modulation of the N400 component also changes in response to the overall statistical structure of the experimental environment: it is modulated less in environments that discourage versus encourage lexico-semantic prediction. For example, in semantic priming paradigms, the degree of N400 attenuation on individual semantically related (versus unrelated) target words is reduced in the presence of a lower (versus a higher) proportion of semantically related word-pairs (Lau, Holcomb, & Kuperberg, 2013; Lau, Weber, Gramfort, Hämäläinen, & Kuperberg, 2014; see also Brown, Hagoort, & Chwilla, 2000; Holcomb, 1988) —a change that is captured by a Bayesian model of expectation adaptation (Delaney-Busch, Lau, Morgan, & Kuperberg, 2017).

Findings like these emphasize the need to consider properties of fillers when deriving predictions for an experiment: if comprehenders readily adapt the strength of their predictions based on the statistics of the recent input, this means that the statistical structure of filler stimuli in an experiment can affect how critical stimuli are processed. We also note that potential sensitivity to the statistics of *all* materials in an experiment does not point to a weakness of prediction as a fundamental mechanism underlying language processing (cf. Ito et al., 2017), but rather to a feature of a system that adapts to changes in the statistics of the input, so as to robustly process noisy perceptual input (for review, Clark, 2013; Kleinschmidt & Jaeger, 2015).

In the specific comparison between DeLong et al. (2005) and Nieuwland et al. (2017), it is not straightforward in what direction the lack of fillers would affect the outcome. On the one hand, some properties of the fillers might lead one to expect *less* N400 modulation on the noun in DeLong et al. (2005), compared to Nieuwland et al. (2017). For example, all fillers in DeLong et al. (2005) contained relatively constraining initial contexts set up by subject-verb combinations (e.g. “Bakers slice….”). Half of the fillers contained highly lexically expected continuations (e.g. “bread”) and half contained less expected continuations (e.g. “pizza”, see DeLong et al., 2017b). For each filler item, highly expected and less expected continuations were counterbalanced across participants. Thus, the relative percentage of sentences in which a relatively strong lexical prediction was confirmed (50%) and disconfirmed (50%) was identical in DeLong et al. (2005) and in Nieuwland et al. (2017). However, before any given experimental trial, participants in DeLong et al. (2005), on average, would have encountered a higher overall *number* of sentences in which a strong lexical prediction was disconfirmed, which may have led them to reduce the degree to which they engaged in predictive processing. On this account, the overall predictability effect on the N400 evoked by nouns would be smaller in DeLong et al. (2005) than in Nieuwland et al. (2017).

On the other hand, other properties of the fillers might lead one to expect *more* N400 modulation on the noun in DeLong et al. (2005), compared to Nieuwland et al. (2017). In DeLong’s (2005) study, the less lexically expected continuations in the fillers were never preceded by an indefinite article. And, in fact, the fillers contained an additional 16 sentences that included indefinite articles, all of which occurred in non-constraining contexts (this is in addition to 24 instances of indefinite articles in non-constraining contexts in the critical items, which appeared in both DeLong et al., 2005 and Nieuwland et al., 2017). The proportion of indefinite *articles* that strongly disconfirmed expectations was thus higher in Nieuwland et al. (2017) than in the original experiment. On this account, the ERP effect on the article would be *smaller* in Nieuwland et al. (2017), compared to DeLong et al.’s (2005) original study.

## The Bayesian analyses

In addition to the analyses we summarized above, Nieuwland et al. (2017) also present two Bayesian analyses. The first analysis builds on Nieuwland and colleagues’ correlation analyses carried out on the datasets in the individual labs (mirroring the analysis performed by DeLong et al. (2005)). However, as discussed above, this analysis did not use baseline-corrected data (differing from DeLong et al., 2005), which might turn out to have made a difference. We therefore do not discuss this analysis further.

The second Bayesian analysis carried out by Nieuwland and colleagues was a trial-level analysis on the full dataset that combined all 9 labs (a Bayesian linear mixed-effects regression). This analysis determined that the credible interval for the effect of cloze probability on the article ranged from [-.06, .69]. Since this credible interval contains zero, Nieuwland et al., argued that this constitutes further evidence for a failure to replicate. However, we note that the majority of credible interval is *larger* than zero, i.e. suggesting an effect of cloze probability on the article in the same direction as that found by DeLong et al. (2005). While this by itself is not strong evidence in favor of the effect reported in DeLong et al. (2005), it is arguably even less expected under the assumption that there is no relation between cloze probability and N400 amplitude on the article (for the same point, see Vasishth, 2017).

## Using cloze probabilities to capture predictability effects

The next two issues that deserve attention relate to the use of cloze probabilities to investigate neural signatures of prediction, including (by hypothesis) the N400. The points we raise here are of relevance not only to the present debate, but to work on neural (and behavioral) signatures of prediction more generally.

### Lack of precision in estimating the cloze probability

Both DeLong et al. (2005) and Nieuwland et al. (2017) operationalize lexical predictability in terms of cloze probabilities. This is, indeed, the most common approach in ERP studies, going back to the original landmark study by Kutas and Hillyard (1984). There are, however, important issues with this approach (for reviews, see Smith & Levy, 2011; Staub, Grant, Astheimer, & Cohen, 2015). These include, for example, potential differences in the type of mechanisms involved in implicit prediction during language processing and those engaged in explicitly filling in blank completions in a production (cloze) task (Smith & Levy, 2011). Here we focus on two other related issues (as also discussed by Smith & Levy, 2013 for reading time analyses). First, the *precision* of cloze probability estimates is a direct function of the number of cloze completions obtained in the norming study. For example, Nieuwland at al. (2017) obtained cloze judgments from 44 participants for the articles, and 30 participants for the nouns. This corresponds to a maximum precision of 1/44 = 0.022 for the articles and 1/30 = 0.033 for the nouns. The same holds for DeLong et al. (2005), who collected data from 30 participants for both articles and nouns. Second, as can be seen in Figure 1, a large proportion of the total test items have a cloze probability of 0 (in DeLong et al., 2005: 30% for the articles, 27.5% for the nouns; in Nieuwland et al., 2017: 20% for the articles, 26.25% for the nouns). But these items do not really have the same predictability, and so, for these items, there is a complete loss of any precision.^3^

Both of these issues—limited precision and complete loss of *any* precision for a large proportion of items (those with cloze probabilities of 0)— are *methodological* concerns. These concerns weigh particularly heavily given that there is now more and more evidence suggesting that behavioral and neural signatures of disconfirmed expectations are primarily driven by small differences *low* predictability continuations, as we discuss next.

### Probability vs. Surprisal

Although the difference in cloze probability between 0 and .022 may seem trivial, it is of concern because some researchers have proposed that the *surprisal* of a word may be a better predictor of processing difficulty than raw probability (Hale, 2001; Levy, 2005). A word’s surprisal—i.e., the logarithm transform of the reciprocal of its probability—is a measure of ‘information’: the amount of information gained upon seeing a word. Because of the logarithmic transform, differences between *small* probabilities close to zero are therefore expected to make a big difference in influencing N400 modulation: the surprisal of a perfectly predictable word with a probability of 1 is 0 bits, and, for each halving in probability, the surprisal doubles. That is, if a response (such as the amplitude of the N400) is linear in surprisal, differences between small probabilities are expected to affect this response more strongly than equal differences between large probabilities. For example, for probabilities of .5 vs. .25, the surprisal is 1 vs. 2 bits, respectively; for probabilities of .015625 vs. .0078125, the surprisal is 6 vs. 7 bits. This means that it is critical to have high precision for low probability events. But that is exactly where cloze probabilities fail.

Empirical support for the claim that surprisal is a better index of probabilistic effects on processing difficulty than raw probability comes primarily from behavioral studies of reading times, which typically have estimated surprisal from language databases (e.g., Smith & Levy, 2013; see also Demberg & Keller, 2008; Frank & Bod, 2011; Linzen & Jaeger, 2016). A few ERP studies have also shown that surprisal predicts the amplitude of the N400 (Delaney-Busch et al., 2017; Frank, Galli, & Vigliocco, 2015; see also Rabovsky, Hansen, & Mcclelland, 2016; Willems, Frank, Nijhof, Hagoort, & Van Den Bosch, 2016). Most of these studies did not directly compared surprisal with raw probability estimates. However, in a recent ERP semantic priming study, Delaney-Busch, Morgan, Lau & Kuperberg (2017) found that word surprisal was a better predictor of N400 amplitude than word probability (Delaney-Busch, Morgan, Lau & Kuperberg, 2017; see also Wlotko & Federmeier, 2012 for further evidence of a non-linear relationship between cloze probability and N400 amplitude).

Based on this literature, it would thus seem that log-transformed cloze probabilities, rather than raw cloze probabilities, might be a better predictor of N400 amplitude. We thus conducted one post-hoc analysis for both the article and the noun using the data generously provided by Nieuwland et al. (2017). We repeated the trial-based analysis (across all nine labs, which used the −100~0ms baseline correction and no filtering, and in which lab was excluded as a predictor), but with log-transformed rather than raw cloze probabilities as the dependent measure. ^4^

We found log-transformed cloze probabilities of the noun to be a significant predictor of N400 amplitude on the noun (p < 1.02 * 10^−14^). This effect size was larger (t =10.34) than for raw cloze probabilities (t = 9.26). Critically, we also found that log-transformed cloze probabilities of the article were a significant predictor of the N400 amplitude on the article (p = 0.015). This latter effect remained significant when we repeated the analysis, including lab both as a fixed effect (p=0.015), and as a random effect with a by-lab intercepts and slopes for cloze effect (p=0.016), and after correcting for the family-wise Type I error rate due to the multiple comparisons.^5^ That is, unlike raw cloze probabilities (reported by Nieuwland et al. 2017), log-transformed cloze probabilities yield an effect on the N400 amplitude on both the noun and the article. We also ran separate regression analyses for each lab separately, again with log-transformed cloze probabilities. As shown in Figure 2, the effect holds in 8 out of the 9 laboratories that participated in Nieuwland et al. (2017).

**Figure 2.**
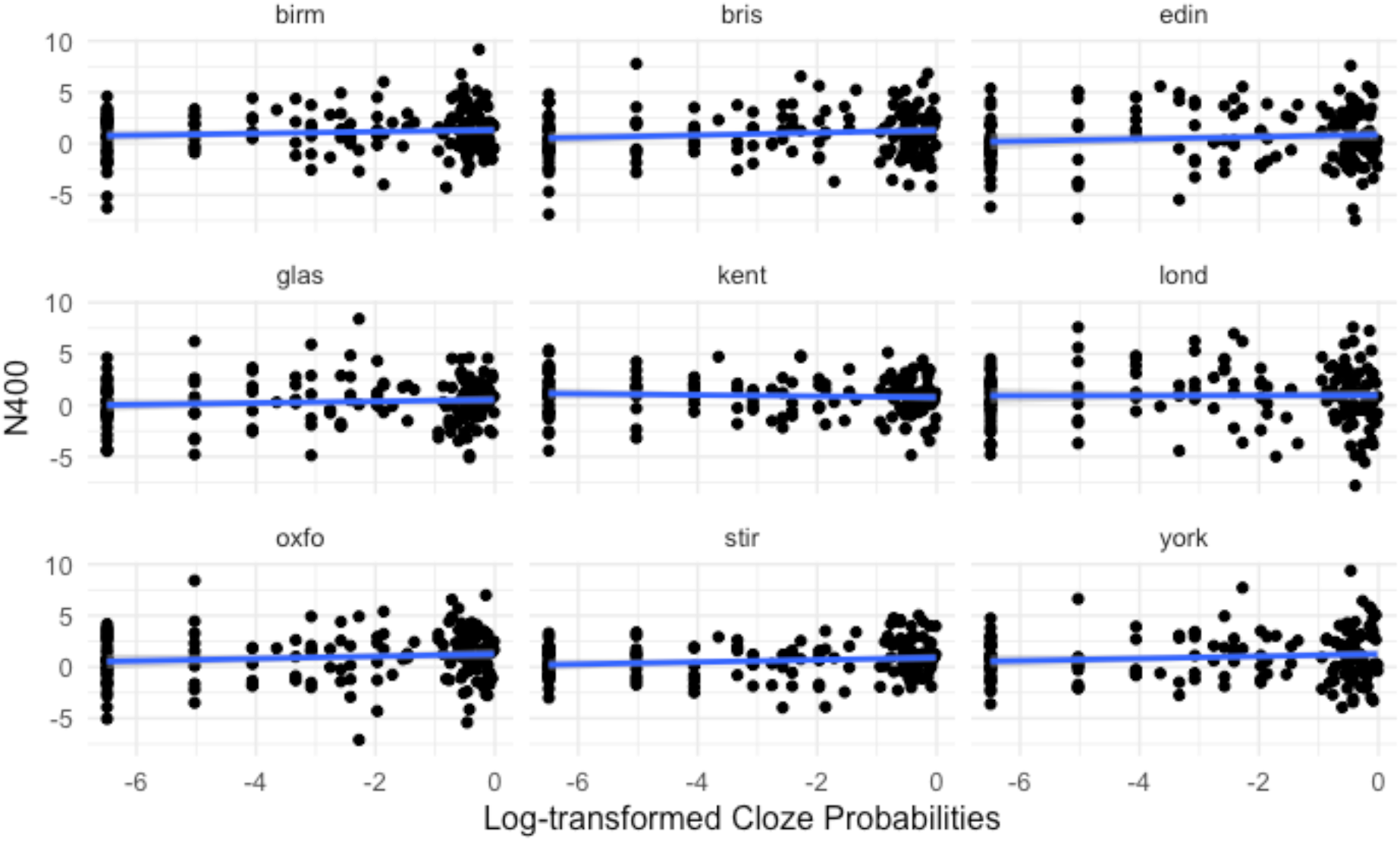
Relationship between log-transformed cloze probabilities and amplitude of N400 on articles for each of the nine labs in Nieuwland et al. (2017).

## Looking beyond 2005

In sum, Nieuwland and colleagues present an impressive effort to replicate an important effect in the field. However, we think that their conclusion that their data provides no evidence for prediction on the article may be premature. First, their design and analyses deviate from those of the original study in ways that might reduce the expected effect size for the N400 evoked at the article. Second, one of the Bayesian analyses would, if anything, seem to support, rather than call into question, the presence of an effect. And third, recent findings suggest that log-transformed cloze probability (or surprisal) should be a better predictor of N400 amplitude. And, indeed, once log-transformed, rather than raw, cloze probabilities are analyzed, we find a significant effect on both the noun *and* the article in 8 out of 9 of the labs that participated in the replication attempt.

In the remainder of this paper, we take a step back and consider the design of DeLong’s original study in relation to the three questions that we posed at the beginning of this paper about the nature of prediction in language processing (see also Kuperberg & Jaeger, 2016). We will argue that a detailed consideration of these issues in relation to DeLong et al.’s design, poses important challenges that future work will need to address. Specifically, we raise questions about how best to assess any effects of prediction—both in relation to the ERP components of interest, and in relation to how indices of probabilistic prediction can be operationalized and formalized. We also discuss different interpretations of the effect that DeLong et al. (2005) report. For the purpose of this discussion, we take the effect that they report at face value, and derive additional predictions that future work (or re-analyses of existing data) could assess.

## What is the timing of prediction in relation to the bottom-up input?

Traditionally, a stark distinction has been drawn between ‘anticipatory processing’ (e.g. DeLong et al., 2005; Kamide, 2008), which is sometimes simply referred to as ‘prediction’, e.g. Federmeier, 2007) and ‘integration’ (Federmeier, 2007; Kamide, 2008; Kuperberg & Jaeger, 2016; Van Petten & Luka, 2012). In this context, *anticipatory processing* refers to the preactivation of relevant information before new bottom-up information becomes available, while *integration* refers to activity at the point at which new bottom-up information becomes available. Of course, ‘integration’ will be influenced by anticipatory activity. Indeed, as pointed out by Kuperberg & Jaeger (2016), it is logically impossible to explain effects of contextual predictability on processing a new input without assuming that the context has already influenced the state of the language processing system prior to this bottom-up input. However, as noted by Nieuwland et al. (2017), effects of context have not always been taken as evidence of anticipatory processing/pre-activation because they can be taken to index “a mixture of attentional and memory retrieval processes [i.e., top-down influences] instigated by the [bottom-up input] itself” (p. 17).

DeLong et al.’s findings on the article have been taken as evidence for anticipatory processing of the predicted noun because they are evident prior to the appearance of this noun. What is not often considered, however, is that the ERP signature being measured at the article is not actually a pure index of anticipatory processing or pre-activation, but rather a reflection of the interaction between such anticipatory activity and activity induced by the appearance of the article itself (this is also true of studies with similar designs in which ERPs are measured to a functional element whose form is dependent on the predicted word, e.g. Van Berkum et al., 2005; Wicha et al., 2004).^6^ This is why any interpretation of what drives differential activity at the article must carefully consider both the type and the level of representation of the information that is being predicted, and what aspects of these predictions become updated when new bottom-up information becomes available. This brings us directly to the next point.

## The nature and the representational level of information that is predicted

Most interpretations of DeLong et al. (2005) seem to assume an account in which any ERP effect at the article is mediated through anticipatory processing of the noun—both its semantic features as well as its phonological properties. We depict one way of elaborating on this position in Figure 3. Here, pre-activation of the noun’s semantic features might stem from prediction at a still higher-level event representation that is based on the comprehender’s current belief about the message that is being conveyed (Kuperberg, 2016; McRae & Matsuki, 2009). This strong semantic pre-activation, in turn, leads to pre-activation of the noun’s phonological form, which, in turn, leads the comprehender to predict the form of the article. As a result, when the less expected article appears, there is increased processing difficulty, leading to differential ERP modulation. What is less often discussed, is the level(s) of representation at which such processing difficulty on the article is incurred. This has direct implications for how we interpret the ERP effect evoked at the article.

**Figure 3.**
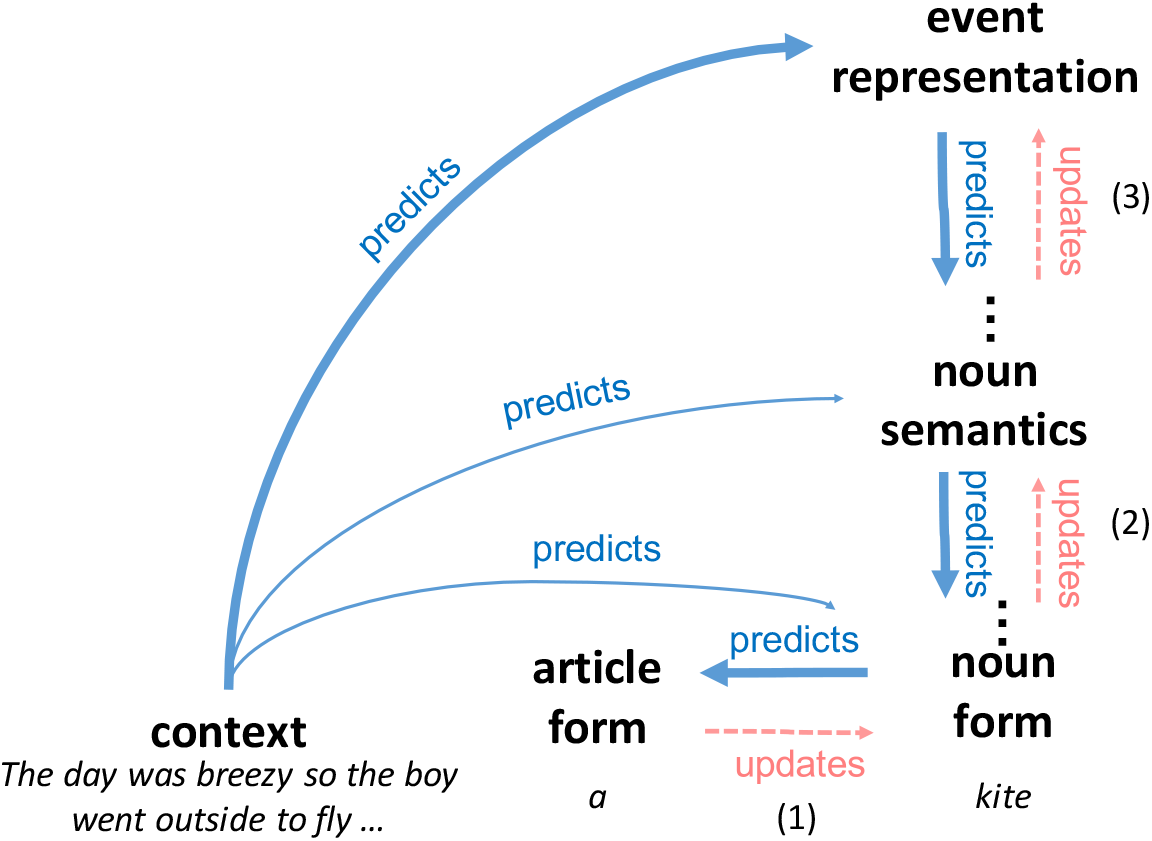
Illustration of a possible predictive chain that would result in differential ERP modulation when the article is encountered. Predictions of the event representation lead to predictions about the semantic properties of the noun (the next referent and head of the next syntactic phrase), which spell out into form predictions of the noun, and form predictions of the article (represented with blue arrows). When the article appears, predictions may be updated at any of these levels (represented with red arrows and labeled by the numbers in parentheses), as outlined in the text.

In theory, an ERP on the article could reflect updating of any or all of the three types of predictions outlined above and as shown in Figure 3: (1) predictions about the form of the article, (2) predictions about the form of the noun, and/or (3) predictions about the semantic features of the noun. Of these three possibilities, (3) would provide the best explanation of why DeLong et al. (2005) reported that the effect on the article had an N400-like temporal and topographic distribution (but see below for discussion of a reanalysis of the same data reported by DeLong, Groppe, Urbach, & Kutas, 2012). Given that the N400 is known to reflect processing at a semantic level (Kutas & Federmeier, 2011), despite being measured on the article, the implicit assumption seems to be that its modulation reflects updating of predictions about semantic properties of the upcoming noun.

However, the focus on the N400 doesn’t rule out the possibility that there may also be earlier effects of updating form predictions (possibilities 1 and 2). In fact, an argument along the lines of Figure 3 *presupposes* form prediction of both the noun (as discussed by both DeLong et al., 2005 and by Nieuwland et al., 2017) as well as of the article. However, the neural indices of any form prediction have not yet been systematically assessed in this particular paradigm. We elaborate on this point below.

Any updating of orthographic or phonological form predictions would likely manifest on ERP components that are distinct from the classic N400, with earlier peaks and possibly different scalp distributions. Several such ERP components have been discussed. These include the N250 (Brothers, Swaab, & Traxler, 2015; Kuperberg, 2013; Lau et al., 2013, see Grainger & Holcomb, 2009 for characterization of the N250 in relation to masked priming paradigms), the phonological mismatch negativity (Connolly & Phillips, 1994) or the N200 (van den Brink, Brown, & Hagoort, 2001) that are associated with phonemic mismatches in spoken language, the P2, which has been linked to the top-down attentionally-mediated extraction of visual features (e.g. Federmeier, Mai, & Kutas, 2005, see also Paczynski & Kuperberg, 2012), and a slightly later positive-going component that has been associated with the confirmation of strong predictions of idioms or frequent collocations and posited to reflect the P300 (Bornkessel-Schlesewsky et al., 2015; Molinaro & Carreiras, 2010; Roehm, Bornkessel-Schlesewsky, Rösler, & Schlesewsky, 2007; Vespignani, Canal, Molinaro, Fonda, & Cacciari, 2009). The peaks of some of these early components are sometimes included within the time windows used to assess the N400 (see Lau et al., 2013, for discussion), and, indeed, their peaks may have been included within the time-window of 200-500ms selected by DeLong et al. (2005) and Nieuwland et al. (2017) to characterize the N400 in their analyses. However, they can be dissociated functionally from the N400, and they sometimes have broader and/or more frontal scalp distributions than the classic centro-parietally distribution associated with the N400. Interestingly, a reanalysis of DeLong’s 2005 data appears in a subsequent paper reported by DeLong et al., (2012: the young controls). Here, in addition to conventional analyses that collapsed across time windows, the authors used mass univariate analyses, which allows for more flexible testing to identify precise time-course and spatial distributions while still maintaining an appropriate Type I error rate (Groppe, Urbach, & Kutas, 2011; Maris & Oostenveld, 2007; see Fields, 2017, for a recent discussion). This analysis showed that the effect on the article did actually have a somewhat different time-course and scalp distribution from the effect observed on the noun: its peak was at medial frontal sites and it was significant between 254ms and 357ms.

Some researchers have also reported contextual predictability effects on the modulation of still earlier components that peak before 200ms, linking this to prediction at the level of early perceptual visual or auditory features. In ERP studies, for example, very early effects have been reported in both studies of reading (e.g., Kim & Lai, 2012; Sereno, Brewer, O’Donnell, & Donnell, 2003), and spoken language comprehension (Groppe et al., 2010). And, in MEG, contextual information can sometimes influence modulation of the visual M1 component evoked by incoming words (Dikker, Rabagliati, Farmer, & Pylkkänen, 2010). Most of the evidence for these very early effects comes from studies in which the input violates very strong structural constraints of the preceding context (Dikker et al., 2010) or very highly semantically constraining contexts (Dikker & Pylkkanen, 2011).^7^ Interestingly, examination of Figure 1 in the original study by DeLong and colleagues suggests some divergence in the waveforms evoked by the high versus low cloze probability articles *before* the 200ms time window. It is possible that, if reliable, this early differences reflects prediction updating of either the article’s or the noun’s visual features. On the other hand, as pointed out by Ito and colleagues (Ito et al., 2017, p. 10-11), some electrode sites in DeLong et al.’s (2005) dataset also show evidence of divergence between the two waveforms *before* the onset of the article (see Figure 2A from DeLong et al., 2012, which shows the same data with a pre-stimulus baseline). Ito et al., (2017) attribute this to “slow signal drift” (also observed in some of the datasets analyzed by Nieuwland et al., 2017, p. 10). Whether this pre-stimulus effect reflects artifact, or whether it reflected a neural correlate of anticipatory processing that manifests even before the onset of the article remains an open question.

Given these arguments, we consider it important that future work directly assesses the critical role of form predictions outlined in Figure 3, and other evidence of form prediction further (although this would not necessarily have to pursued within the DeLong et al., paradigm). Because the earlier ERP components linked to form processing are less well characterized than the N400, one cannot simply collapse across *a priori* time windows and electrode sites to test relevant hypotheses. However, the use of mass univariate analyses provides a potentially powerful alternative way of testing relevant hypotheses while still correcting for multiple comparisons (Groppe et al., 2011; Maris & Oostenveld, 2007). More generally, we consider it critical to spell out specific hypotheses about the nature of the predictive chains hypothesized to underlie effects of context in the ERP signal.^8^

## Is prediction probablistic in nature?

DeLong et al.’s study was designed to test the hypothesis that anticipatory processing, as reflected by ERP modulation at the point of the article, was probabilistic and graded in nature. The precise level and nature of the processing actually reflected by ERP modulation on the article is not spelled out. However, as discussed above, a common interpretation of the N400-like effect reported by DeLong and colleagues is that it reflects anticipatory *semantic* processing. The gradient measure of prediction strength that DeLong et al. (2005)—and, thus also Nieuwland et al. (2017)—employ to test this hypothesis is the *probability of the article itself* (which, when log-transformed, translates on to the surprisal of the article). To the best of our knowledge, neither Delong and colleagues, nor subsequent work, has considered the question of *why* the probability of the article itself would be a good index of degree to which *semantic* predictions of the upcoming noun are updated during the processing of the article. As we show next, a principled probabilistic measure of the degree of semantic prediction updating on the article differs from— but is correlated with—the surprisal of the article.

In the language of probability theory, updates of predictions upon receiving new information can be formalized as the degree to which comprehenders change their beliefs. In the present case, this should be a function of the objective shift in the probability distribution over all noun semantics from before seeing the article to after seeing the article (see Figure 4).

**Figure 4.**
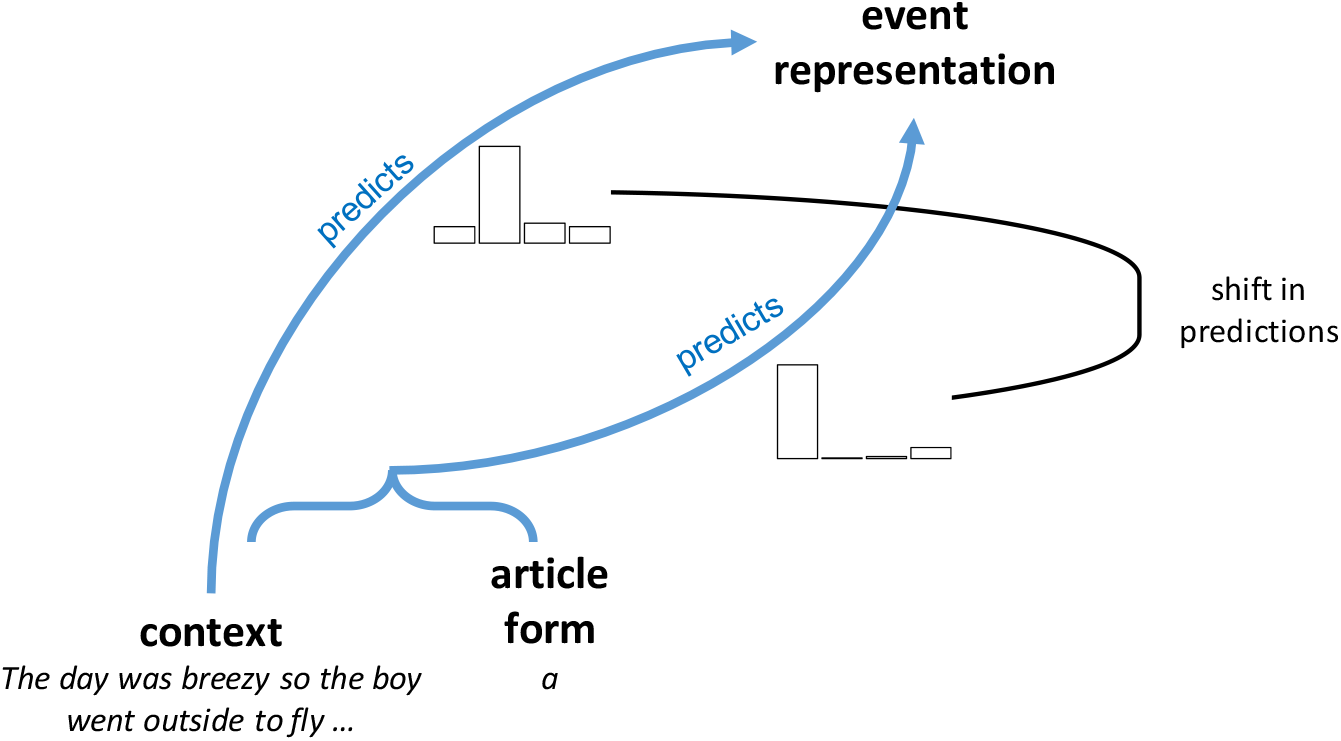
Predictions about upcoming noun shift upon observing the article.

One principled probabilistic measure of this shift in predictions about the upcoming noun semantics from before to after encountering the article, is the relative entropy or Kullback-Leibler divergence between the distribution over all possible upcoming noun semantics before and after seeing the article. This is also known as *Bayesian surprise* (Doya, Ishii, Pouget, & Rao, 2007; Itti & Baldi, 2009). For example, after seeing the article ‘a’, the Bayesian surprise about the upcoming noun semantics (NOUNi) can be expressed as:^9^

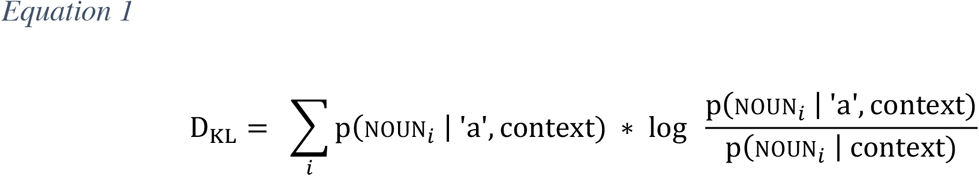

Bayesian surprise has been used successfully to, for example, model the cost of incremental argument integration (Hörberg, 2016), and recent simulations using a neural network model link Bayesian surprise to the pattern of N400 modulation during sentence comprehension (Rabovsky et al., 2016). However, to the best of our knowledge, no analysis so far has estimated the effect of Bayesian surprise on the N400-like response on the article in DeLong et al.’s (2005) paradigm. Upon encountering the article, ‘a’, the predicted probability of each possible noun semantics changes from p(noun_i_ | context) to p(noun_i_ | ‘a’, context), where the latter can be re-expressed as:

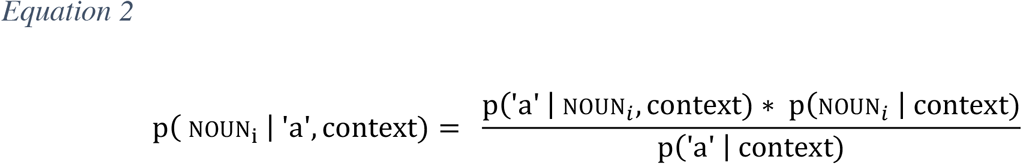

Inserting (E2) back into (E1) to calculate the Bayesian surprise (over the distribution of noun semantics) incurred while processing the article, we get the (E3):

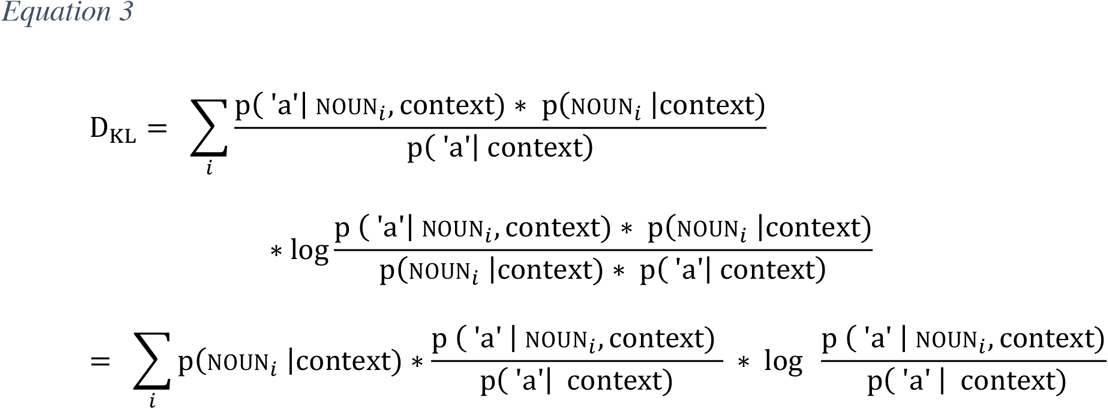

which can be further reorganized (for details, see appendix):

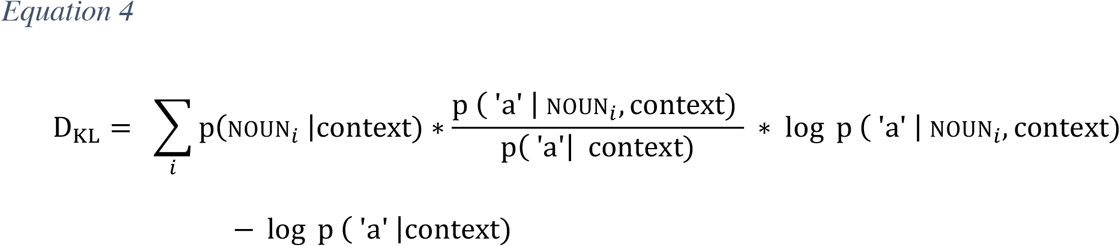

That is, the Bayesian surprise incurred on the article is the sum of the article’s surprisal, −log p (‘a’ |context), plus another component, the first term in (E4). The Bayesian surprise over the noun semantics thus differs from both the probability of the article itself, p (‘a’ | context), and the article’s surprisal, log 1 / p ( ‘a’ | context). Unlike the article’s surprisal, the Bayesian surprise also depends on probability of article form given each predicted noun semantics, p (‘a’ | noun_*i*_, context). The two indices are, however, closely related. First, since log p (‘a’ | noun_*i*_, context) is always smaller than or equal to zero, the first term in (E4) will always be negative. The Bayesian surprise over the noun semantics on the article will thus never be larger than the surprisal of the article. Second, for contexts in which the article depends deterministically on the noun, i.e., p (‘a’ | noun_*i*_, context) always equals either 1 or 0, the Bayesian surprise over the noun semantics reduces to the article’s surprisal. That is, for those types of contexts, D_KL_ = − log p (‘a’ | context) (for proof, see appendix).^10^

Such a deterministic relation is, however, unlikely to hold for the stimuli in DeLong et al. (2005) and Nieuwland et al. (2017): the form of the English indefinite article is determined by the word that immediately follows it, and that word is not *always* the noun (Nieuwland and colleagues estimate the probability of a noun occurring directly after an article to be only .33; our own estimates from different corpora of American English ranged from ~.3 in writing to ~.7 in speech). This raises the question how the article’s surprisal is related to the Bayesian surprise over the noun semantics in the data sets of DeLong et al. (2005) and Nieuwland et al. (2017).

We therefore explored the relation between the two probabilistic indices based on a simple language model and data from natural language use.^11^ Specifically, we extracted all non-sentence initial noun phrases from three corpora of British and American English. We set the context in (E3) to the one word immediately preceding the noun phrase to reduce data sparsity. Table 1 lists the number of context-noun combinations and the distribution of the frequencies of these combinations for each corpus. For each context, we calculated the surprisal of ‘a’ (and ‘an’) as well as the Bayesian surprise over the distribution of nouns on ‘a’ (and ‘an’). As shown in Figure 5, these two probabilistic indices are highly correlated.

**Table 1.** Characteristics of different copora adopted in the analysis

| Corpus Name | Citation | Variety of English | Genre | Mode | #Words | # One-word Contexts | log<sub>10</sub> Context Frequency Mean (SD) |
| --- | --- | --- | --- | --- | --- | --- | --- |
| Switchboard | Marcus, Santorini, Marcinkiewicz, & Taylor, 1999 | American | Conversation | Spoken | ~800,000 | 660 | 1.28 (0.60) |
| Brown |  | American | Mixed | Written | ~1 million | 736 | 1.16 (0.54) |
| Wall Street Journal |  | American | Newspaper | Written | ~2 million | 1869 | 1.18 (0.50) |
| British National Corpus | British National Corpus Consortium, 2007 | British | Mixed | Written | ~90 million | 20943 | 1.47 (0.70) |

**Figure 5.**
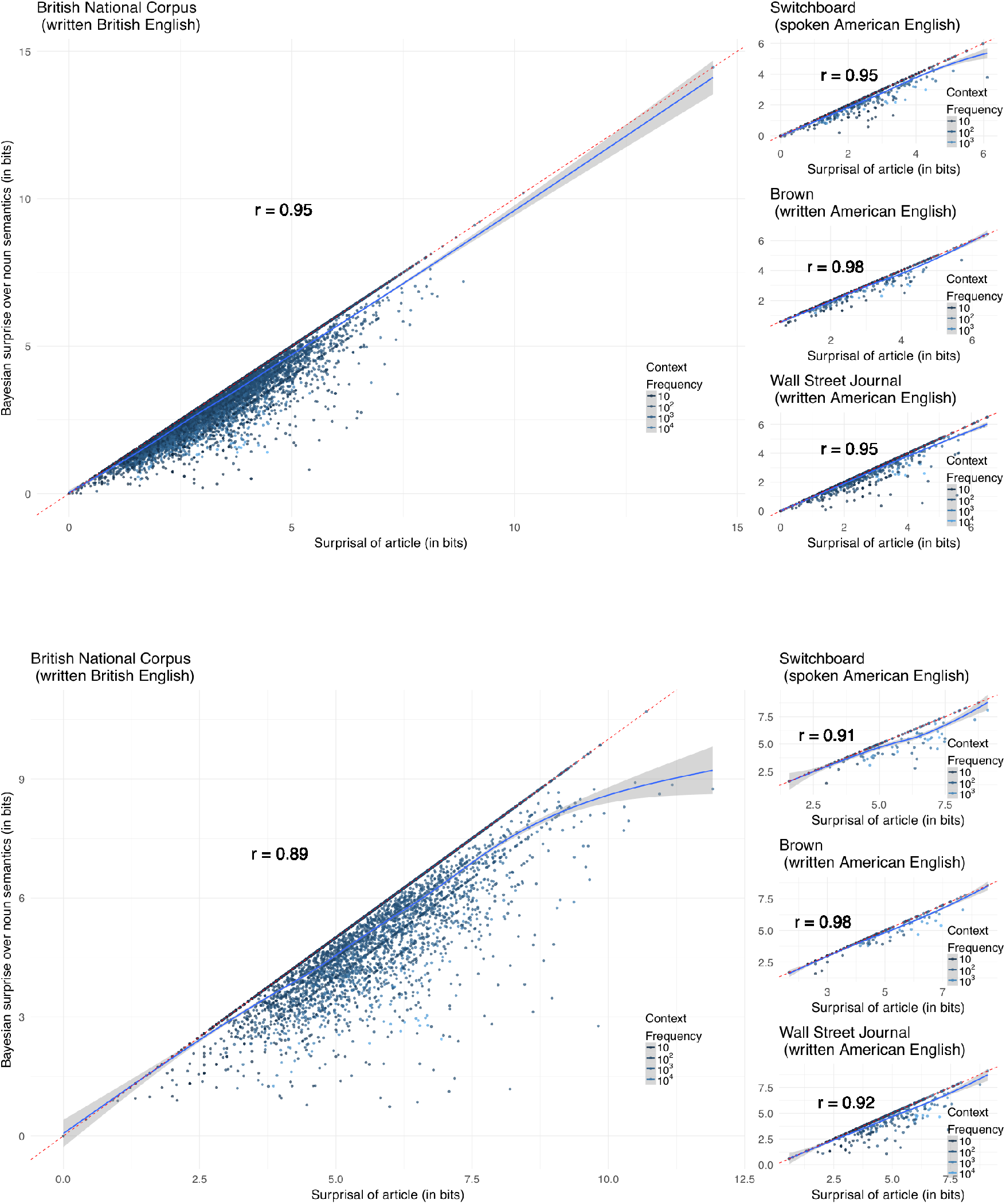
Correlation between the article’s surprisal and the Bayesian surprise over the distribution of the upcoming noun incurred on the article. Both indices were estimated for four separate corpora. For details, see text. Each dot represents a context immediately preceding a noun phrase. Blue line shows non-parametric smoother predicting Bayesian surprise from surprisal. Top: data for ‘a’. Bottom: data for ‘an’.

**Figure 6.**
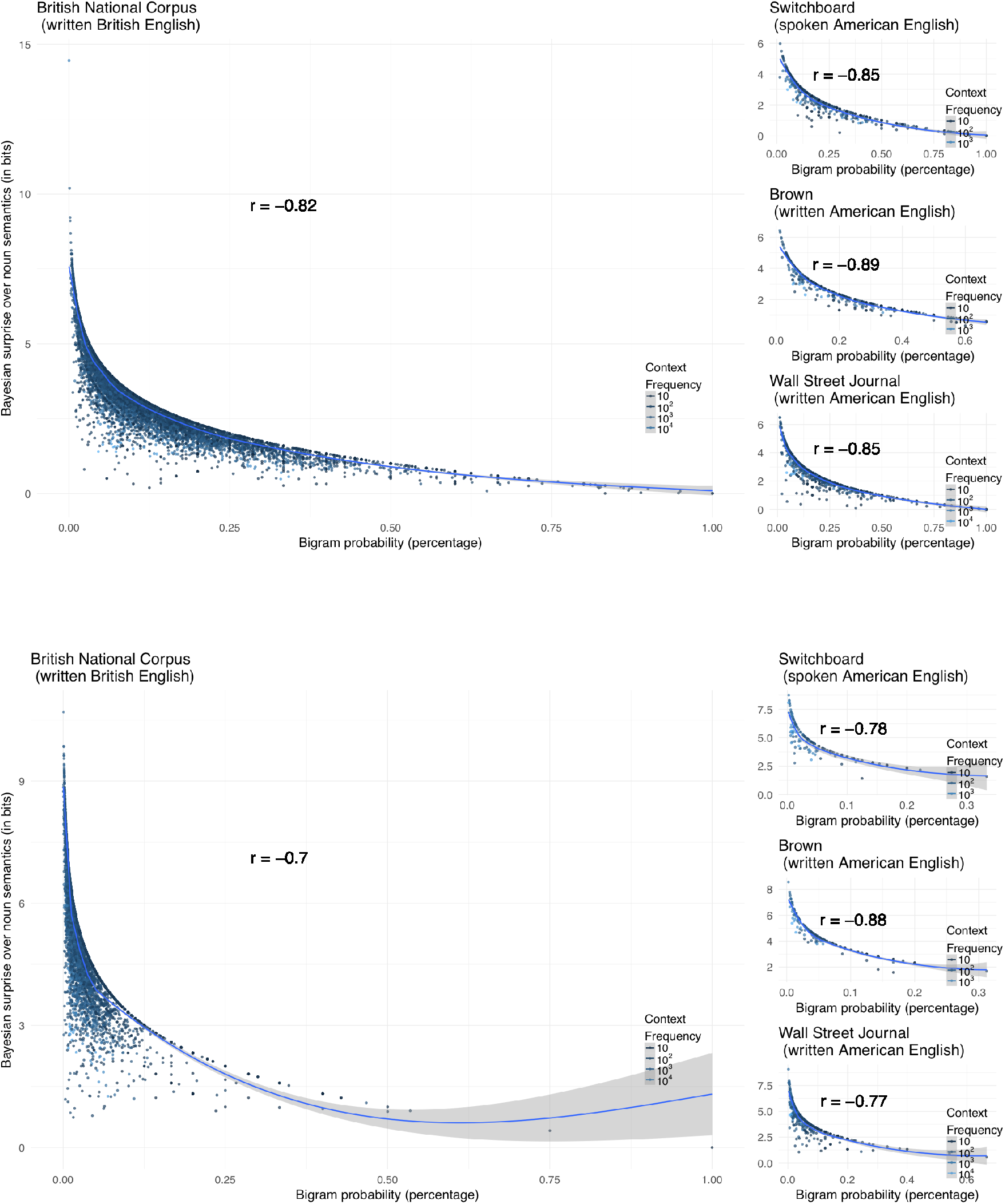
Correlation between the article’s predictability (bi-gram probability) and the Bayesian surprise over the distribution of the upcoming noun incurred on the article. Both indices were estimated for four separate corpora. For details, see text. Each dot represents a context immediately preceding a noun phrase. Blue line shows non-parametric smoother predicting Bayesian surprise from article predictability. Top: data for ‘a’. Bottom: data for ‘an’.

We emphasize that our estimates of the two indices are crude, potentially over-estimating the correlation between them. We also emphasize that here we calculated the Kullback-Leibler divergence over noun *forms* from before having seen the article (i.e., based on the one-word preceding context) to after having seen the article. With the types of stimuli used by DeLong et al. (2005), the distribution of noun forms approximates the distribution of noun semantics. This assumption, however, is not always appropriate, as the predictability of a given word-form can be dissociated from the predictability of its semantic features. And, critically, the amplitude of the N400 is primarily sensitive to the latter (see also Kuperberg, 2016 for recent discussion). Still, to the extent that similarly high correlations between this measure of Bayesian surprise, and surprisal of the article hold for the stimuli in DeLong et al. (2005) and Nieuwland et al. (2017), this would provide an explanation for why the article’s surprisal correlated with the N400—a neural response that is typically associated with *semantic* processing.

When one inspects the relationship between the predictability of the article (as shown in Figure 6), it is clear that Bayesian surprise does not have a linear relationship with article predictability. Despite so, Bayesian surprise is also correlated with the predictability of the article, although the correlation is weaker than that with the surprisal of the article. This might explain why, although article predictability is not directly related to N400 amplitude, when DeLong et al. (2005) found correlation between N400 amplitude and article predictability and Nieuwland et al. (2017) found a trend in the same direction in their by-trial level analysis.

## Summary

Regardless of questions about its replicability, the findings of DeLong et al. (2005) inspire new ways of thinking about prediction. For the goals of the original study, when there was still debate about whether or not anticipatory semantic processing occurred at all during language comprehension, it was arguably not critical to determine the exact nature of the computational processes reflected by the ERP component evoked by the article. Rather, the data presented by DeLong et al. (2005), along with that of other studies employing similar designs (e.g. Van Berkum et al., 2005; Wicha et al., 2004) was important in that it provided evidence that probabilistic anticipatory processing happened at all.

Since this original publication, however, the field has advanced and the questions that need to be asked next are of a more specific nature. The point of discussion presented above is to illustrate the value of a careful consideration of the nature of prediction in guiding hypotheses and interpretation. We believe that it is important for future work to distinguish between predictive processes at different levels of representation. The strongest tests of such processes will commit to testing precise relationships between probabilistic indices and (sets of) ERP components that are thought to reflect processing at the level of representation linked to the hypothesized predictive process. Specifically, we have argued that, so long as any N400 modulation on the article is interpreted as reflecting updating of predictions about the noun’s semantic properties, then Bayesian surprise (reflecting shifts in predictions about the upcoming noun from before to after the article is encountered) is a more appropriate probabilistic index than the cloze probability of the article. While we have also shown that the article’s *surprisal* provides an easy approximation of Bayesian surprise over the noun’s semantics incurred on the article, the two indices are not the same either (cf. Figure 5), and future studies might be able to tease them apart.

If one is interested in whether comprehenders are updating their predictions about the *form* features of the noun, then Bayesian surprise would also be the predictor of interest. In this case, however, it would be more appropriate to examine its relationship with the modulation of earlier ERP components that are associated with form processing, and that have a different temporal and topographic distribution than the N400 effect, as discussed above. If, however, one is interested in updating of predictions about the form of the article (rather than noun), then the cloze probability or surprisal of the article is the most principled predictor, and, once again, it would be more appropriate to examine its relationship with earlier ERP components.

## Conclusions

In summary, we think that it is premature to conclude that that Nieuwland et al.’s dataset provides no evidence for anticipatory processing at the point of the article. In this paper, we have discussed several differences between the impressive replication effort reported in Nieuwland et al. (2017) and the original study by DeLong and colleagues. Further, our own re-analysis of the data obtained by Nieuwland and colleagues finds a significant correlation between the amplitude of the N400 on the article and the article’s surprisal (rather than its raw cloze probability). The article’s surprisal would be a principled probabilistic index of predictions of the article’s form. We also showed that the article’s surprisal is very strongly correlated with a principled probabilistic index of the updating of semantic predictions about the upcoming noun (the Bayesian surprise or Kullback-Leibler divergence).

On the other hand, we do echo Nieuwland and colleagues’ emphasis on the importance of future studies probing prediction at the level of phonological form. As discussed above, the current findings leave open the question of whether the ERP effects observed by DeLong and colleagues (2005, see also DeLong et al., 2012), as well as the effects revealed by our re-analysis of Nieuwland et al. (2017), should be attributed to updating at the level of form or semantic features (or both), and whether these effects will withstand further scrutiny. Indeed, the question of whether form prediction exists, and what role in plays in language comprehension, continues to be a matter of great interest. Beyond language processing, some have called into question that higher-level expectation *ever* spells out into low-level form prediction (cf. Firestone & Scholl, 2015 and responses to it). Within research on spoken language processing, there is a lively debate about the existence of feedback during word recognition. While some models assume feedback from lexical representations to pre-lexical processes (Elman & McClelland, 1988; Magnuson, McMurray, Tanenhaus, & Aslin, 2003; McClelland & Elman, 1986), others have argued that existing evidence can be explained without reference to feedback (McQueen, Cutler, & Norris, 2003; Norris, McQueen, & Cutler, 2000; Norris & McQueen, 2008; Norris, McQueen, & Cutler, 2015); for discussion, see also Kuperberg & Jaeger, 2016). While either view is compatible with *some* types of form prediction (e.g., form prediction based on phonological context, McQueen et al., 2003), the latter explicitly argues against influences of lexical—and thus semantic—influences on early form processing.

More generally, these issues bear upon the two extreme positions that Nieuwland et al. (2017) argue against in their paper. The first is that we always “pre-activate words at all levels of representation in a routine and implicit (i.e., non-strategic) fashion”, and the second is that prediction down to lexical form is necessarily associated with the prediction-through-language-production (Pickering & Garrod, 2013). Regarding the first claim, this seems, to us, somewhat of a strawman. Like DeLong et al. (2017b), we are not aware of anyone who has made the claim that we necessarily or always predict at every level of representation during comprehension. Regarding the second claim, prediction at the level of form does not necessarily imply that the production system is used to generate predictions during comprehension. Rather, we think that it is more likely that both comprehenders and producers draw upon common generative models, where *generative* neither implies nor rules out the involvement of production circuits (Brown & Kuperberg, 2015; Jaeger & Ferreira, 2013; Kleinschmidt & Jaeger, 2015).

The real question at stake, we submit, is not *whether* we can engage in probabilistic prediction, but to what degree and under what circumstances, and to what level(s) of representation (see Kuperberg & Jaeger, 2016 for discussion). This recent debate between Nieuwland and colleagues and DeLong and colleagues offers a welcome opportunity to revisit and clarify these questions, which is a necessary step towards understanding the nature of predictive language comprehension.

## Acknowledgments

We thank all the labs who participated in the 9-lab study for contributing to this important enterprise and for sharing their data. We applaud Nieuwland et al. for coordinating the study and sharing the preprocessed data and code, and leading by example in pre-registering their study. We are grateful to Katherine DeLong and Mante Nieuwland for patiently answering our many questions about their approach and data, and for sharing with us the raw data from their cloze task as well as the set of fillers they used. We are also very grateful to Nate Delaney-Busch, Albert Kim, Andrea Martin-Nieuwland, and Mante Nieuwland for insightful discussions surrounding this topic. This work was partially supported by NICHD R01 HD082527 to GRK and NIH R01 HD075797 as well as NSF CAREER IIS-1150028 to TFJ. The views expressed here are those of the authors and do not necessarily reflect views of the funding agencies.

## Appendix

### Appendix A Bayesian surprise and article surprisal

In this appendix, we derive the relations between the article’s surprisal and the Bayesian surprise over the noun semantics on the article described in the main text. To get from (E8) to (E9):

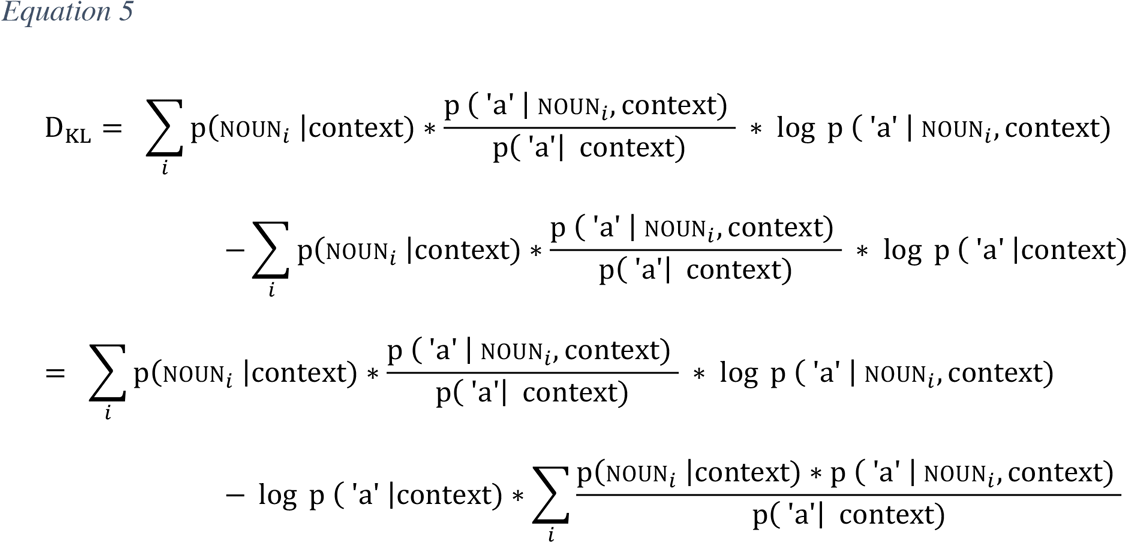

Since Σ_*i*_ p(noun_*i*_ |context) * p (‘a’ | noun_*i*_, context) = p(‘a’ |context), the second term can be further simplified, yielding:

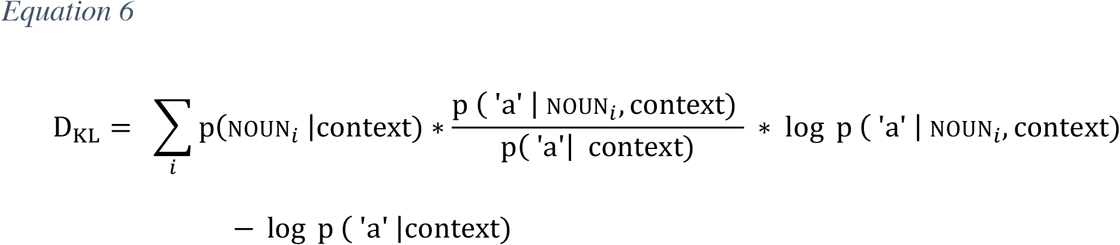

To show that this reduces to the article’s surprisal, when the article depends deterministically on the noun, i.e., p (‘a’ | noun_*i*_, context) always equals either 1 or 0:

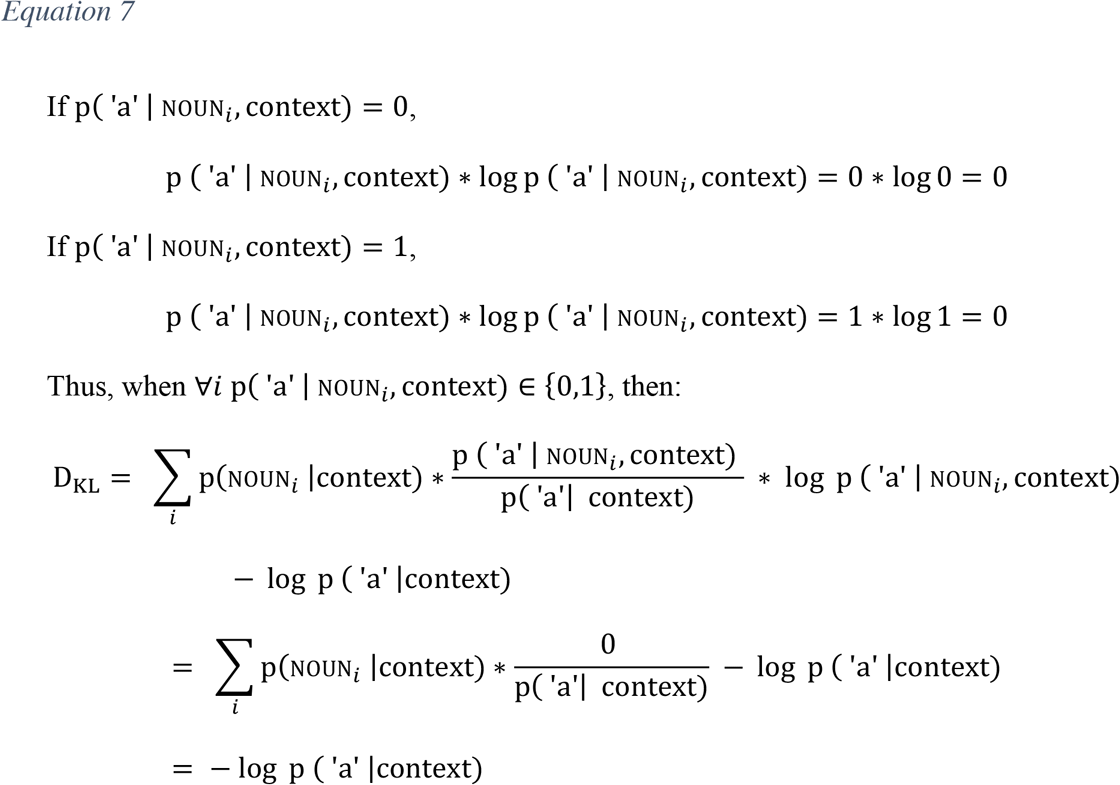

### Appendix B Bayesian surprise and article surprisal for (certain) agreement systems

In some languages with agreement, the form of prenominal articles is determined by the noun regardless of whether the noun immediately follows the article. In those environments, the surprisal of the article is identical to the Bayesian surprise with regard to the noun, for the reasons outlined in appendix A. For example, when the predicted noun is masculine, the probability of encounter a masculine article p(masc | noun_*i*_, context) will always equal 1. Hence we can derive the Bayesian surprise upon seeing a masculine article as:

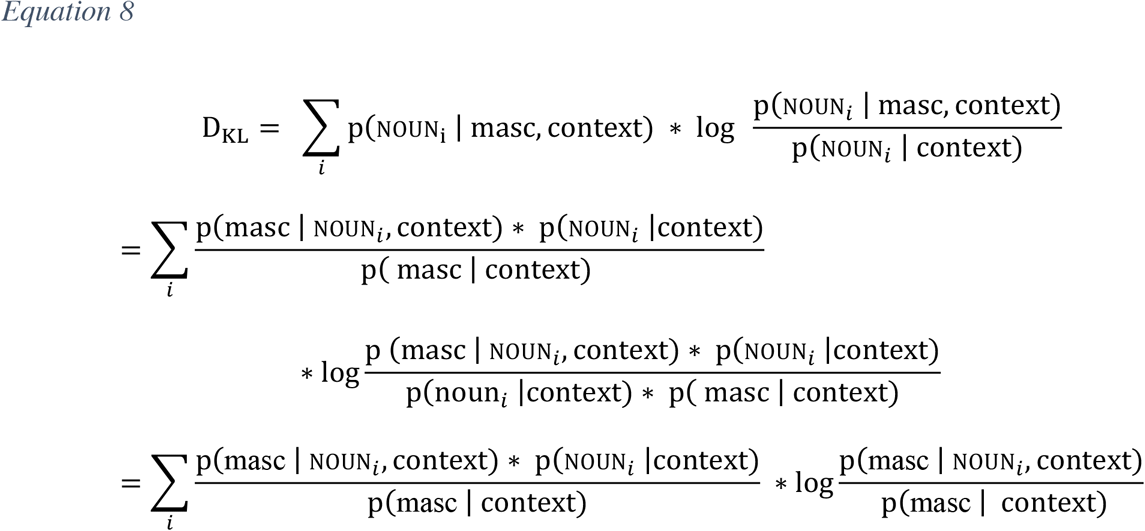

Since the gender agreement between the article and the noun is not affected by intervening elements, p( masc | noun_*i*_, context) equals one if noun_i_ is masculine, and equals zero if noun_i_ is not masculine. Therefore the Bayesian surprise can be further simplified as:

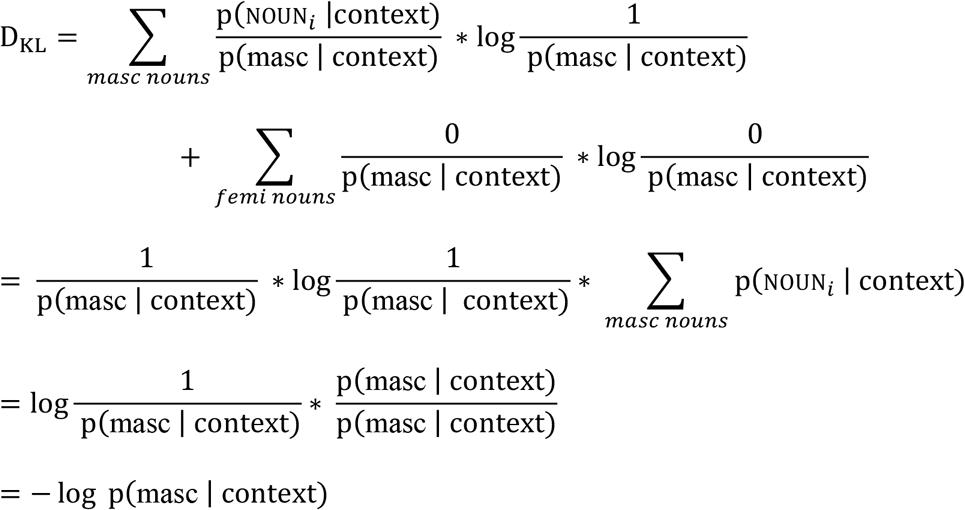

For such systems, a correlation between neural responses and the cloze probability (or surprisal) of gender-marked articles could thus reflect updating of prediction about the noun’s semantics, predictions about the article form, (non-anticipatory) processing of the semantics inherent in the gender-marking itself (as pointed out in Nieuwland et al., 2017), or any combination of these. Interestingly, studies on predictive processing in these types of agreement systems have found a variety of neural signatures (e.g., Otten et al., 2007; Otten & Van Berkum, 2007, 2008; Van Berkum et al., 2005; Wicha et al., 2004; for discussion, see Nieuwland et al., 2017). One possible explanation of this is that stimulus- and design-specific differences between these studies lead to differences in the extent to which these different possibilities are reflected in the ERPs.

## Footnotes

1 We note that, while the rule that determines the form of the indefinite article is phonological (‘an’ before vowel onsets, ‘a’ in all other contexts), there is a close correspondence to orthography (but see, e.g., ‘a uniform’, which phonologically starts with a consonant, but orthographically with a vowel character). Since words violating this correspondence were not included in the experiment matrials used by DeLong et al. (2005), the relevant form prediction could be orthographic or phonological or both.

2 Nieuwland et al. (2017) also conduct additional single trial analyses of the N400 amplitude on the article using different prestimulus baseline corrections (−200~0ms and −500~0ms). With the −200~0ms baseline correction, the N400 analysis on the article still trends in the expected direction (p<.19); with the, −500~0ms baseline correction, p > .50. They also used a high-pass filer of .1Hz combined with a −100~0ms baseline correction, and the effect on article is non-significant (p > .56).

3 This issue is hard to overcome with cloze tasks: the number of participants required to obtain reliable larger-than-zero counts for low probability words would often be in the order of thousands or tens of thousands.

4 Log-transforming cloze probabilities requires smoothing to deal with items with 0 cloze probabilities. We follow a standard approach to this smoothing problem (plus 1 smoothing with a uniform prior). A better solution to be considered in future studies is to elicit cloze judgments from many more participants, so as to avoid/reduce this problem altogether.

5 Log cloze probabilities are also a significant predictor for −200~0ms baseline-corrected N400 amplitudes (p = 0.047), and show the same numerical trend when the −500~0ms baseline was used (p = 0.277) and when the −100~0ms baseline was used in combination with 0.1 high-pass filtering (p = 0.15). We note that the analysis reported in the main text was the first we performed. It corresponds to the approach advocated by Nieuwland and colleagues.

6 When DeLong et al. (2005) was first published, there was already evidence from the visual world paradigm for anticipatory processing, which was not dependent on measuring activity that was time-locked to new bottom-up input. This evidence for anticipatory processing has been argued by some (Nieuwland et al., 2017; Huettig & Mani, 2016) to critically depend on the restricted set of potential candidates that is typical to visual world studies (for evidence against this interpretation, see Dahan, Magnuson, & Tanenhaus, 2001; for discussion, see Salverda & Tanenhaus, 2017). Since DeLong’s initial publication, additional evidence for anticipatory processing has come from studies reporting differential neural modulation that is not time-locked to new bottom-up input (e.g., Boylan et al., 2014; Dikker & Pylkkänen, 2013; Piai et al., 2016; Sohoglu, Peelle, Carlyon, & Davis, 2012).

7 We note that such early effects need to be interpreted with caution: early ERP and MEG components are highly sensitive to differences across conditions in bottom-up perceptual inputs, and even when these features are tightly counterbalanced across conditions, differences may arise as a result of excluding trials with artifact during preprocessing, leading to the spurious early effects.

8 In this context, we note that it is possible that form predictions of the article proceed directly on the basis of contextual information without being mediated through the noun semantics (e.g., a direct gradient association between *ngram* or latent semantic presentations of the context with a specific form of the article). As the indefinite article is generally assumed not to evoke event or semantic processing, an argument along these lines would have to explain the presense of a N400-like response — e.g., as reflecting a later signature of form prediction (see above).

9 The equations we provide describe the updating of predictions about the upcoming noun semantics. We are not considering uncertainty about *at what point in the sentence* the noun (form or syntactic category) will appear. This strikes us as a reasonable assumption given that predictions can be assumed to be generated on the basis of the message-level (or similar) representation.

10 In the appendix we also show that, for this very reason, Bayesian surprise completely reduces to the article’s surprisal for certain agreement phenomena for which anticipatory processing has also been studied (Van Berkum et al., 2005; Wicha et al., 2004). For those agreement systems, the noun *does* deterministically affect the form of the article.

11 An alternative would be to estimate the relation between the two indices specifically for the stimuli in the experiment, based on the raw cloze responses. However, this approach suffers from data sparsity for reasons related to those raised in our discussion of cloze norms above (this was confirmed after annotating the raw cloze reponses from DeLong et al., 2005).

## References

Allopenna, P. D., Magnuson, J. S., & Tanenhaus, M. K. (1998). Tracking the Time Course of Spoken Word Recognition Using Eye Movements: Evidence for Continuous Mapping Models. Journal of Memory and Language, 38(38), 419–439. https://doi.org/10.1006/jmla.1997.2558

Altmann, G. T. M., & Kamide, Y. (2007). The real-time mediation of visual attention by language and world knowledge: Linking anticipatory (and other) eye movements to linguistic processing. Journal of Memory and Language, 57(4), 502–518. https://doi.org/10.1016/j.jml.2006.12.004

Bornkessel-Schlesewsky, I., Philipp, M., Alday, P. M., Kretzschmar, F., Grewe, T., Gumpert, M., … Schlesewsky, M. (2015). Age-related changes in predictive capacity versus internal model adaptability: Electrophysiological evidence that individual differences outweigh effects of age. Frontiers in Aging Neuroscience, 7, 217. https://doi.org/10.3389/fnagi.2015.00217

Boylan, C., Trueswell, J. C., & Thompson-Schill, S. L. (2014). Multi-voxel pattern analysis of noun and verb differences in ventral temporal cortex. Brain and Language, 137, 40–49. https://doi.org/10.1016/j.bandl.2014.07.009

British National Corpus Consortium. (2007). British National Corpus version 3 (BNC XML edition). Oxford: Oxford University Computing Services.

Brothers, T., Swaab, T. Y., & Traxler, M. J. (2015). Effects of prediction and contextual support on lexical processing: Prediction takes precedence. Cognition, 136, 135–149. https://doi.org/10.1016/j.cognition.2014.10.017

Brown-Schmidt, S. (2009). Partner-specific interpretation of maintained referential precedents during interactive dialog. Journal of Memory and Language, 61(2), 171–190. https://doi.org/10.1016/j.jml.2009.04.003

Brown, C. M., Hagoort, P., & Chwilla, D. J. (2000). An event-related brain potential analysis of visual word priming effects. Brain and Language, 72(2), 158–190. https://doi.org/10.1006/brln.1999.2284

Brown, M., & Kuperberg, G. R. (2015). A Hierarchical Generative Framework of Language Processing: Linking Language Perception, Interpretation, and Production Abnormalities in Schizophrenia. Front. Hum. Neurosci, 9, 1–23. https://doi.org/10.3389/fnhum.2015.00643

Connolly, J. F., & Phillips, N. A. (1994). Event-Related Potential Components Reflect Phonological and Semantic Processing of the Terminal Word of Spoken Sentences. Journal of Cognitive Neuroscience, 6(3), 256–266. https://doi.org/10.1162/jocn.1994.6.3.256

Creel, S. C., Aslin, R. N., & Tanenhaus, M. K. (2008). Heeding the voice of experience: The role of talker variation in lexical access. Cognition, 106(2), 633–664. https://doi.org/10.1016/j.cognition.2007.03.013

Dahan, D., Magnuson, J. S., & Tanenhaus, M. K. (2001). Time Course of Frequency Effects in Spoken-Word Recognition: Evidence from Eye Movements. Cognitive Psychology, 367, 317–367. https://doi.org/10.1006/cogp.2001.0750

Dahan, D., & Tanenhaus, M. K. (2004). Continuous mapping from sound to meaning in spoken-language comprehension: immediate effects of verb-based thematic constraints. Journal of Experimental Psychology. Learning, Memory, and Cognition, 30(2), 498–513. https://doi.org/10.1037/0278-7393.30.2.498

Delaney-Busch, N., Lau, E. F., Morgan, E., & Kuperberg, G. R. (2017). Comprehenders Rationally Adapt Semantic Predictions to the Statistics of the Environment: a Bayesian Model of trial-level N400 amplitudes. In Proceedings of the 39th Annual Conference of the Cognitive Science Society. London, UK.

DeLong, K. A., Groppe, D. M., Urbach, T. P., & Kutas, M. (2012). Thinking ahead or not? Natural aging and anticipation during reading. Brain and Language, 121(3), 226–239. https://doi.org/10.1016/j.bandl.2012.02.006

Delong, K. A., Urbach, T. P., & Kutas, M. (2017). Concerns with Nieuwland et al. (2017). Retrieved from http://kutaslab.ucsd.edu/FinalDUK17Comment9LabStudy.pdf

DeLong, K. A., Urbach, T. P., & Kutas, M. (2005). Probabilistic word pre-activation during language comprehension inferred from electrical brain activity. Nature Neuroscience, 8(8), 1117–1121. https://doi.org/10.1038/nn1504

DeLong, K. A., Urbach, T. P., & Kutas, M. (2017). Is there a replication crisis? Perhaps. Is this an example? No: A commentary on Ito, Martin & Nieuwland (2016). Language, Cognition, and Neuroscience. Advance online publication. https://doi.org/10.1080/23273798.2017.1279339

Demberg, V., & Keller, F. (2008). Data from eye-tracking corpora as evidence for theories of syntactic processing complexity. Cognition, 109(2), 193–210. https://doi.org/10.1016/j.cognition.2008.07.008

Dikker, S., & Pylkkanen, L. (2011). Before the N400: Effects of lexical-semantic violations in visual cortex. Brain and Language, 118(1–2), 23–28. https://doi.org/10.1016/j.bandl.2011.02.006

Dikker, S., & Pylkkänen, L. (2013). Predicting language: MEG evidence for lexical preactivation. Brain and Language, 127(1), 55–64. https://doi.org/10.1016/j.bandl.2012.08.004

Dikker, S., Rabagliati, H., Farmer, T. A., & Pylkkänen, L. (2010). Early occipital sensitivity to syntactic category is based on form typicality. Psychological Science, 21(5), 629–34. https://doi.org/10.1177/0956797610367751

Doya, K., Ishii, S., Pouget, A., & Rao, R. P. N. (Eds.). (2007). Bayesian brain: Probabilistic approaches to neural coding. MIT Press. https://doi.org/10.7551/mitpress/9780262042383.001.0001

Elman, J. L., & McClelland, J. L. (1988). Cognitive penetration of the mechanisms of perception: Compensation for coarticulation of lexically restored phonemes. Journal of Memory and Language, 27(2), 143–165. https://doi.org/10.1016/0749-596X(88)90071-X

Farmer, T. A., Fine, A. B., Yan, S., & Cheimariou, S. (2014). Error-Driven Adaptation of Higher-Level Expectations During Reading. In Proceedings of the 36th Annual Meeting of the Cognitive Science Society (Vol. 1, pp. 2181–2186).

Federmeier, K. D. (2007). Thinking ahead: the role and roots of prediction in language comprehension. Psychophysiology, 44(4), 491–505. https://doi.org/10.1111/j.1469-8986.2007.00531.x

Federmeier, K. D., Mai, H., & Kutas, M. (2005). Both sides get the point: hemispheric sensitivities to sentential constraint. Memory & Cognition, 33(5), 871–886. https://doi.org/10.3758/BF03193082

Fields, E. C. (2017). Event-related Potential and Functional MRI Studies of Emotion and Self Relevance. Tufts University.

Fine, A. B., & Jaeger, T. F. (2016). The Role of Verb Repetition in Cumulative Structural Priming in Comprehension. Journal of Experimental Psychology: Learning Memory and Cognition, 42(9), 1362–1376. https://doi.org/10.1037/xlm0000236

Fine, A. B., Jaeger, T. F., Farmer, T. A., & Qian, T. (2013). Rapid expectation adaptation during syntactic comprehension. PLoS ONE, 8(10). https://doi.org/10.1371/journal.pone.0077661

Fine, A. B., Qian, T., Jaeger, T. F., & Jacobs, R. A. (2010). Is there syntactic adaptation in language comprehension? In Proceedings of ACL-2010: Workshop on Cognitive Modeling and Computational Linguistics (pp. 18–26).

Firestone, C., & Scholl, B. J. (2015). Cognition does not affect perception: Evaluating the evidence for “top-down” effects. Behavioral and Brain Sciences, 4629, 1–77. https://doi.org/10.1017/S0140525X15000965

Frank, S. L., & Bod, R. (2011). Insensitivity of the human sentence-processing system to hierarchical structure. Psychological Science, 22(6), 829–834. https://doi.org/10.1177/0956797611409589

Frank, S. L., Otten, L. J., Galli, G., & Vigliocco, G. (2015). The ERP response to the amount of information conveyed by words in sentences. Brain and Language, 140, 1–11. https://doi.org/10.1016/j.bandl.2014.10.006

Fraundorf, S. H., & Jaeger, T. F. (2016). Readers generalize adaptation to newly-encountered dialectal structures to other unfamiliar structures. Journal of Memory and Language, 91, 28–58. https://doi.org/10.1016/j.jml.2016.05.006

Grainger, J., & Holcomb, P. J. (2009). Watching the word go by: On the time-course of component processes in visual word recognition. Linguistics and Language Compass, 3(1), 128–156. https://doi.org/10.1111/j.1749-818X.2008.00121.x

Groppe, D. M., Choi, M., Huang, T., Schilz, J., Topkins, B., Urbach, T. P., & Kutas, M. (2010). The phonemic restoration effect reveals pre-N400 effect of supportive sentence context in speech perception. Brain Research, 1361, 54–66. https://doi.org/10.1016/j.brainres.2010.09.003

Groppe, D. M., Urbach, T. P., & Kutas, M. (2011). Mass univariate analysis of event-related brain potentials/fields I: A critical tutorial review. Psychophysiology, 48(12), 1711–1725. https://doi.org/10.1111/j.1469-8986.2011.01273.x

Hale, J. (2001). A probabilistic earley parser as a psycholinguistic model. In Proceedings of the Second Meeting of the North American Chapter of the Association for Computational Linguistics on Language Technologies (pp. 159–166). https://doi.org/doi:10.3115/1073336.1073357

Holcomb, P. J. (1988). Automatic and attentional processing: An event-related brain potential analysis of semantic priming. Brain and Language, 35(1), 66–85. https://doi.org/10.1016/0093-934X(88)90101-0

Hörberg, T. (2016). Probabilistic and Prominence-driven Incremental Argument Interpretation in Swedish. Stockholm University.

Huettig, F., & Mani, N. (2015). Is prediction necessary to understand language? Probably not. Language, Cognition, and Neuroscience, 3798, 1–26. https://doi.org/10.1080/23273798.2015.1072223

Ito, A., Martin, A. E., & Nieuwland, M. S. (2016). How robust are prediction effects in language comprehension? Failure to replicate article-elicited N400 effects. Language, Cognition and Neuroscience. Advance online publication. https://doi.org/10.1080/23273798.2016.1242761

Ito, A., Martin, A. E., & Nieuwland, M. S. (2017). Why the a/an prediction effect might be hard to replicate: A rebuttal to DeLong, Urbach & Kutas (2017). Language, Cognition and Neuroscience. Advance online publication. doi:10.1080/23273798.2017.1323112.

Itti, L., & Baldi, P. (2009). Bayesian surprise attracts human attention. Vision Research, 49(10), 1295–1306. https://doi.org/10.1016/j.visres.2008.09.007

Jaeger, T. F., & Ferreira, V. S. (2013). Seeking predictions from a predictive framework. The Behavioral and Brain Sciences, 36(4), 359–60. https://doi.org/10.1017/S0140525X12002762

Kamide, Y. (2008). Anticipatory Processes in Sentence Processing. Linguistics and Language Compass, 2(4), 647–670. https://doi.org/10.1111/j.1749-818X.2008.00072.x

Kaschak, M. P. (2007). Long-Term Structural Priming Affects Subsequent Patterns of Language Production. Memory & Cognition, 35(5), 925–937. https://doi.org/10.3758/BF03193466

Kaschak, M. P., & Glenberg, A. M. (2004). This construction needs learned. Journal of Experimental Psychology. General, 133(3), 450–467. https://doi.org/10.1037/0096-3445.133.3.450

Kim, A., & Lai, V. (2012). Rapid interactions between lexical semantic and word form analysis during word recognition in context: evidence from ERPs. Journal of Cognitive Neuroscience, 24(5), 1104–12. https://doi.org/10.1162/jocn_a_00148

Kleinschmidt, D. F., Fine, A. B., & Jaeger, T. F. (2012). A belief-updating model of adaptation and cue combination in syntactic comprehension. Proceedings of the 34th Annual Meeting of the Cognitive Science Society (CogSci12), 599–604.

Kleinschmidt, D. F., & Jaeger, T. F. (2015). Robust speech perception: Recognize the familiar, generalize to the similar, and adapt to the novel. Psychological Review, 122(2), 148–203. https://doi.org/10.1037/a0038695

Kuperberg, G. R. (2013). The Proactive Comprehender: What Event-Related Potentials tell us about the dynamics of reading comprehension. In Unraveling the Behavioral, Neurobiological, and Genetic Components of Reading Comprehension (pp. 1–23). Baltimore: Paul Brookes Publishing.

Kuperberg, G. R. (2016). Separate streams or probabilistic inference? What the N400 can tell us about the comprehension of events. Language, Cognition and Neuroscience, 31(5), 602–616. https://doi.org/10.1080/23273798.2015.1130233

Kuperberg, G. R., & Jaeger, T. F. (2016). What do we mean by prediction in language comprehension? Language Cognition & Neuroscience, 31(1), 32–59. https://doi.org/10.1080/23273798.2015.1102299

Kutas, M., DeLong, K. A., & Smith, N. J. (2011). A Look around at What Lies Ahead: Prediction and Predictability in Language Processing. In Predictions in the Brain: Using Our Past to Generate a Future. https://doi.org/10.1093/acprof:oso/9780195395518.003.0065

Kutas, M., & Federmeier, K. D. (2011). Thirty years and counting: Finding meaning in the N400 component of the event related brain potential (ERP). Annual Review of Psychology, 62, 621. https://doi.org/10.1146/annurev.psych.093008.131123

Kutas, M., & Hillyard, S. (1984). Brain potential during reading reflect word expentancy and semantic association. Nature, 307, 161–163.

Lau, E. F., Holcomb, P. J., & Kuperberg, G. R. (2013). Dissociating N400 effects of prediction from association in single-word contexts. Journal of Cognitive Neuroscience, 25(3), 484–502. https://doi.org/10.1162/jocn_a_00328

Lau, E. F., Weber, K., Gramfort, A., H??m??l??inen, M. S., & Kuperberg, G. R. (2014). Spatiotemporal Signatures of Lexical-Semantic Prediction. Cerebral Cortex, 26(4), 1377–1387. https://doi.org/10.1093/cercor/bhu219

Levy, R. (2005). Probabilistic Models of Word Order and Syantactic Discontinuity. Stanford University.

Magnuson, J. S., McMurray, B., Tanenhaus, M. K., & Aslin, R. N. (2003). Lexical effects on compensation for coarticulation: the ghost of Christmash past. Cognitive Science, 27, 285–298.

Marcus, M., Santorini, B., Marcinkiewicz, M. A., & Taylor, A. (1999). Treebank-3 LDC99T42. In Linguistic Data Consortium. Philadelphia.

Maris, E., & Oostenveld, R. (2007). Nonparametric statistical testing of EEG- and MEG-data. Journal of Neuroscience Methods, 164(1), 177–190. https://doi.org/10.1016/j.jneumeth.2007.03.024

McClelland, J. L., & Elman, J. L. (1986). The TRACE model of speech perception. Cognitive Psychology, 18(1), 1–86. https://doi.org/10.1016/0010-0285(86)90015-0

McQueen, J. M., Cutler, A., & Norris, D. (2003). Flow of information in the spoken word recognition system. Speech Communication, 41, 257–270. https://doi.org/10.1016/S0167-6393(02)00108-5

McRae, K., & Matsuki, K. (2009). People use their knowledge of common events to understand language, and do so as quickly as possible. Linguistics and Language Compass, 3(6), 1417–1429. https://doi.org/10.1111/j.1749-818X.2009.00174.x

Molinaro, N., & Carreiras, M. (2010). Electrophysiological evidence of interaction between contextual expectation and semantic integration during the processing of collocations. Biological Psychology, 83(3), 176–190. https://doi.org/10.1016/j.biopsycho.2009.12.006

Nieuwland, M. S., Politzer-Ahles, S., Heyselaar, E., Segaert, K., Darley, E., Kazanina, N., … Huettig, F. (2017). Limits on prediction in language comprehension: A multi-lab failure to replicate evidence for probabilistic pre-activation of phonology. https://doi.org/10.1101/111807

Norris, D., & McQueen, J. M. (2008). Shortlist B: A Bayesian Model of Continuous Speech Recognition. Psychological Review, 115(2), 357–395. https://doi.org/10.1037/0033-295X.115.2.357

Norris, D., McQueen, J. M., & Cutler, A. (2000). Merging information in speech recognition: feedback is never necessary. The Behavioral and Brain Sciences, 23(3), 299–325; https://doi.org/10.1017/S0140525X00003241

Norris, D., McQueen, J. M., & Cutler, A. (2015). Prediction, Bayesian inference and feedback in speech recognition. Language, Cognition and Neuroscience, 3798, 1–15. https://doi.org/10.1080/23273798.2015.1081703

Otten, M., Nieuwland, M. S., & Van Berkum, J. J. A. (2007). Great expectations: Specific lexical anticipation influences the processing of spoken language. BMC Neuroscience, 8, 89. https://doi.org/10.1186/1471-2202-8-89

Otten, M., & Van Berkum, J. J. A. (2007). What makes a discourse constraining? Comparing the effects of discourse message and scenario fit on the discourse-dependent N400 effect. Brain Research, 1153(1), 166–177. https://doi.org/10.1016/j.brainres.2007.03.058

Otten, M., & Van Berkum, J. J. A. (2008). Discourse-Based Word Anticipation During Language Processing: Prediction or Priming? Discourse Processes, 45(6), 464–496. https://doi.org/10.1080/01638530802356463

Paczynski, M., & Kuperberg, G. R. (2012). Multiple influences of semantic memory on sentence processing: Distinct effects of semantic relatedness on violations of real-world event/state knowledge and animacy selection restrictions. Journal of Memory and Language, 67(4), 426–448. https://doi.org/10.1016/j.jml.2012.07.003

Piai, V., Anderson, K. L., Lin, J. J., Dewar, C., Parvizi, J., Dronkers, N. F., & Knight, R. T. (2016). Direct brain recordings reveal hippocampal rhythm underpinnings of language processing. Proceedings of the National Academy of Sciences, 113(40), 1136611371. https://doi.org/10.1073/pnas.1603312113

Pickering, M. J., & Garrod, S. (2013). An integrated theory of language production and comprehension. Behavioral and Brain Sciences, 1–19. https://doi.org/10.1017/S0140525X12001495

Rabovsky, M., Hansen, S. S., & Mcclelland, J. L. (2016). N400 Amplitudes Reflect Change in a Probabilistic Representation of Meaning: Evidence From a Connectionist Model. In A. Papafragou, D. Grodner, D. Mirman, & J. C. Trueswell (Eds.), Proceedings of the 38th Annual Conference of the Cognitive Science Society. Austin, TX: Cognitive Science Society.

Roehm, D., Bornkessel-Schlesewsky, I., Rösler, F., & Schlesewsky, M. (2007). To predict or not to predict: influences of task and strategy on the processing of semantic relations. Journal of Cognitive Neuroscience, 19(8), 1259–1274. https://doi.org/10.1162/jocn.2007.19.8.1259

Salverda, A. P., & Tanenhaus, M. K. (2017). The Visual World Paradigm. In A. M. B. de Groot & P. Hagoort (Eds.), Research methods in psycholinguistics: A practical guide. Malden, MA: Wiley-Blackwell.

Sereno, S. C., Brewer, C. C., & O’Donnell, P. J. (2003). Context Effects in Word Recognition. Psychological Science, 14(4), 328–333. https://doi.org/10.1111/1467-9280.14471

Smith, N. J., & Levy, R. (2011). Cloze but no cigar: The complex relationship between cloze, corpus, and subjective probabilities in language processing. In Proceedings of the 33rd Annual Meeting of the Cognitive Science Conference (pp. 1637–1642).

Smith, N. J., & Levy, R. (2013). The effect of word predictability on reading time is logarithmic. Cognition, 128(3), 302–319. https://doi.org/10.1016/j.cognition.2013.02.013

Sohoglu, E., Peelle, J. E., Carlyon, R. P., & Davis, M. H. (2012). Predictive Top-Down Integration of Prior Knowledge during Speech Perception. Journal of Neuroscience, 32(25), 8443–8453. https://doi.org/10.1523/JNEUROSCI.5069-11.2012

Staub, A., Grant, M., Astheimer, L., & Cohen, A. (2015). The influence of cloze probability and item constraint on cloze task response time. Journal of Memory and Language, 82, 1–17. https://doi.org/10.1016/j.jml.2015.02.004

Van Berkum, J. J. A., Brown, C. M., Zwitserlood, P., Kooijman, V., & Hagoort, P. (2005). Anticipating upcoming words in discourse: evidence from ERPs and reading times. Journal of Experimental Psychology. Learning, Memory, and Cognition, 31(3), 443–467. https://doi.org/10.1037/0278-7393.31.3.443

van den Brink, D., Brown, C. M., & Hagoort, P. (2001). Electrophysiological evidence for early contextual influences during spoken-word recognition: N200 versus N400 effects. Journal of Cognitive Neuroscience, 13(7), 967–985. https://doi.org/10.1162/089892901753165872

Van Petten, C., & Luka, B. J. (2012). Prediction during language comprehension: Benefits, costs, and ERP components. International Journal of Psychophysiology, 83(2), 176–190. https://doi.org/10.1016/j.ijpsycho.2011.09.015

Vasishth, S. (2017). Nieuwland et al replication attempts of DeLong at al. 2005. Retrieved July 4, 2017, from http://vasishth-statistics.blogspot.de/2017/04/a-comment-on-delong-et-al-2005-nine.html

Vespignani, F., Canal, P., Molinaro, N., Fonda, S., & Cacciari, C. (2009). Predictive Mechanisms in Idiom Comprehension. Journal of Cognitive Neuroscience, 22(8), 1682–1700.

Wicha, N. Y. Y., Moreno, E. M., & Kutas, M. (2004). Anticipating Words and Their Gender: An Event-related Brain Potential Study of Semantic Integration, Gender Expectancy, and Gender Agreement in Spanish Sentence Reading. Cognitive Neuroscience, 16(7), 1272–1288. https://doi.org/10.1162/0898929041920487.Anticipating

Willems, R. M., Frank, S. L., Nijhof, A. D., Hagoort, P., & Van Den Bosch, A. (2016). Prediction during Natural Language Comprehension. Cerebral Cortex, 26(6), 2506–2516. https://doi.org/10.1093/cercor/bhv075

Wlotko, E. W., & Federmeier, K. D. (2012). Age-related Changes in the Impact of Contextual Strength on Multiple Aspects of Sentence Comprehension. Psychophysiology, 49(6), 770–785.

Yan, S., & Farmer, T. A. (2015). Adaptation to Unexpected Word-Forms in Highly Predictive Sentential Contexts. In Proceedings of the 28th Annual Meeting of the CUNY Conference on Human Sentence Processing. Los Angeles, CA.

Yildirim, I., Degen, J., Tanenhaus, M. K., & Jaeger, T. F. (2016). Talker-specificity and adaptation in quantifier interpretation. Journal of Memory and Language, 87, 128–143. https://doi.org/10.1016/j.jml.2015.08.003

